# Identifying key underlying regulatory networks and predicting targets of orphan C/D box *SNORD116* snoRNAs in Prader-Willi syndrome

**DOI:** 10.1101/2023.10.03.560773

**Authors:** Rachel B. Gilmore, Yaling Liu, Christopher E. Stoddard, Michael S. Chung, Gordon G. Carmichael, Justin Cotney

**Author notes:** To whom correspondence should be addressed: Dr. Justin Cotney.

## Abstract

Prader-Willi syndrome (PWS) is a rare neurodevelopmental disorder characterized principally by initial symptoms of neonatal hypotonia and failure-to-thrive in infancy, followed by hyperphagia and obesity. It is well established that PWS is caused by loss of paternal expression of the imprinted region on chromosome 15q11-q13. While most PWS cases exhibit megabase-scale deletions of the paternal chromosome 15q11-q13 allele, several PWS patients have been identified harboring a much smaller deletion encompassing primarily *SNORD116*. This finding suggests *SNORD116* is a direct driver of PWS phenotypes. The *SNORD116* gene cluster is composed of 30 copies of individual *SNORD116* C/D box small nucleolar RNAs (snoRNAs). Many C/D box snoRNAs have been shown to guide chemical modifications of other RNA molecules, often ribosomal RNA (rRNA). However, *SNORD116* snoRNAs are termed ‘orphans’ because no verified targets have been identified and their sequences show no significant complementarity to rRNA. It is crucial to identify the targets and functions of *SNORD116* snoRNAs because all reported PWS cases lack their expression. To address this, we engineered two different deletions modelling PWS in two distinct human embryonic stem cell (hESC) lines to control for effects of genetic background. Utilizing an inducible expression system enabled quick, reproducible differentiation of these lines into neurons. Systematic comparisons of neuronal gene expression across deletion types and genetic backgrounds revealed a novel list of 42 consistently dysregulated genes. Employing the recently described computational tool snoGloBe, we discovered these dysregulated genes are significantly enriched for predicted *SNORD116* targeting versus multiple control analyses. Importantly, our results showed it is critical to use multiple isogenic cell line pairs, as this eliminated many spuriously differentially expressed genes. Our results indicate a novel gene regulatory network controlled by *SNORD116* is likely perturbed in PWS patients.

## Introduction

Prader-Willi syndrome (PWS [OMIM #176270]) is a rare, neurodevelopmental disorder characterized by neonatal hypotonia and failure-to-thrive during infancy, followed by hyperphagia and obesity; small stature, hands, and feet; mild to moderate cognitive deficit; and a range of behavioral and sleep problems (Cassidy & Driscoll, 2009; Holm et al., 1993; Prader et al., 1956). PWS is linked to instability of chromosome 15 at locus 15q11-q13 that can result in inheritance of a variety of chromosomal structural changes (Glenn et al., 1997; S. J. Kim et al., 2012). The most common structural change in PWS patients is the loss of several megabases of the 15q11-13 locus specifically on the paternally inherited allele. This is linked to the fact that many of the genes in this region are imprinted, a phenomenon in which genes are expressed exclusively from one parental allele. This imprint is established in the germline via DNA methylation on the maternal allele at the Prader-Willi Syndrome Imprinting Center (PWS-IC) (Brannan I & Bartolomei, 1999; Nicholls et al., 1998; Shemer et al., 2000). The PWS-IC is a promoter for a complex transcriptional unit that includes protein-coding genes (SNURF and SNRPN), many species of small nucleolar RNAs (*SNORD65*, *SNORD108*, two copies of *SNORD109*, 30 copies of *SNORD116,* and 48 copies of *SNORD115*), antisense RNA that can silence *UBE3A* (*UBE3A-ATS*), and other RNA species that are not well understood (Ariyanfar & Good, 2022; Cavaillé et al., 2000; Gray et al., 1999; Rougeulle et al., 1998; Runte et al., 2001). In addition to DNA methylation at the PWS-IC, post-translational methylation modifications have also been found in this chromosomal region. Zinc finger protein ZNF274 has been found to bind specifically to SNORD116 DNA sequences (Langouët et al., 2018, 2020). This binding event is thought to recruit lysine methyltransferases SETDB1 and EHMT2 (Cruvinel et al., 2014; Y. Kim et al., 2017) which results in deposition of methylation marks on lysine 9 of histone H3 (H3K9me3), an epigenetic mark frequently associated with heterochromatin and gene silencing. There are other protein coding genes that are also imprinted as this locus, *MKRN3*, *MAGEL2* and *NDN*, but are positioned upstream of the PWS-IC and governed by different promoter sites. Notably, mutations in *MAGEL2* cause Schaaf-Yang syndrome (SYS [OMIM #615547]), another rare neurodevelopmental disorder which shares some phenotypes with PWS (Schaaf & Marbach, 2021).

While megabase-scale deletions are the most common genetic subtype of PWS, a handful of patients have been reported to have atypical microdeletions (Tan et al., 2020). These deletions specifically effect the tandem array of 30 copies of *SNORD116. SNORD116* is a member of the C/D box class of small nucleolar RNAs (snoRNAs). *SNORD116* can be further subdivided into three subgroups based on sequence similarity: Group I (*SNOG1*, *SNORD116-1* to *SNORD116-9*), Group II (*SNOG2*, *SNORD116-10* to *SNORD116-24*), and Group III (*SNOG3*, *SNORD116-25* to *SNORD116-30*)(Castle et al., 2010; Runte et al., 2001). snoRNAs are generally thought to be processed by exonucleolytic trimming from the introns of a host gene (Tamás Kiss & Filipowicz, 1995) and serve as a scaffold and specificity factor for ribonucleoprotein complexes that deposit 2’-O methylation on maturing ribosomal RNAs (rRNAs)(Filipowicz et al., 1999). However, *SNORD116,* as well as the other snoRNAs found in the 15q11-13 region, do not have sequence complementarity to rRNA. Thus, it is unclear if they participate in rRNA maturation and are typically referred to as orphans (T. Kiss, 2001). A previous study utilized the BLASTn algorithm to predict *SNORD116* sites transcriptome wide (Baldini et al., 2022). However, only a handful of predicted targets were interrogated in HeLa cells making it unclear if they are relevant for PWS.

Since the function of *SNORD116* thus far has remained elusive, much effort has recently been expended to identify gene expression patterns that are dysregulated in PWS. Several studies have compared gene expression between tissue or cell lines derived from PWS patients and those from unrelated controls (Bochukova et al., 2018; Falaleeva et al., 2015; Huang et al., 2021; Sledziowska et al., 2023; Victor et al., 2021). While each of these studies identified numerous genes with distinct expression patterns in the PWS context, a coherent set of consistently dysregulated disease relevant genes has not been identified. Inherent differences in genetic background or postmortem delay may obscure important gene expression changes, leading to lack of a consensus set of perturbed genes in the disorder. Therefore, we have turned to the use of isogenic human embryonic stem cell (hESC) lines, to provide a more rigorous approach to investigate cellular deficits in disease models. Here, we describe the generation of two distinct hESC lines, each engineered with two separate deletions relevant to determining the targets and functions of *SNORD116* snoRNAs. We also utilized an inducible Neurogenin-2 (NGN2) expression system to enable quick, reproducible differentiation of these lines into neurons (Fernandopulle et al., 2018). Performing bulk RNA-sequencing on resulting neurons allowed us to identify a novel list of 42 genes consistently transcriptionally dysregulated in our PWS-like systems. Importantly, our results showed it is critical to use multiple isogenic cell line pairs as this eliminated many spuriously differentially expressed genes. Employing the recently described computational tool snoGloBe (Deschamps-Francoeur et al., 2022), we discovered these dysregulated genes are significantly enriched for predicted *SNORD116* targeting versus multiple control analyses. Our results indicate a novel gene regulatory network controlled by *SNORD116* is likely perturbed in PWS patients.

## Results

### Isogenic cell line pairs utilizing an inducible neuron system were generated to evaluate the effects of *SNORD116* loss in the context of PWS

We initially set out to identify genes that might be consistently dysregulated in PWS. Several studies have reported differentially expressed genes (DEGs) between postmortem brain tissue and iPSC-derived neurons from PWS patients and controls. However, when we analyzed these differential gene expression data, few genes were consistently dysregulated in the disease context (Supplemental Figure 1)(Bochukova et al., 2018; Huang et al., 2021). Further, the genes that were shared between these studies do not show clear connections to PWS-related phenotypes through gene ontology analysis (Supplemental Figure 1). We reasoned that one major contributor to this lack of concordance could be due to differences in genetic backgrounds between PWS patients and controls. To generate models of PWS that could be directly compared to isogenic controls, we engineered two different deletions on the paternal chromosome 15q allele in two distinct hESC lines by utilizing CRISPR/Cas9 editing with guide RNAs (gRNAs) designed to target up- and downstream of our regions of interest (Methods)(Supplemental Table 1). One deletion spanned from alternative promoters of the *SNRPN* transcript upstream of the PWS-IC to the distal end of the *SNORD116* snoRNA cluster (termed “lgDEL” model). The other deletion encompassed just the *SNORD116* cluster (termed “smDEL” model). All six cell lines (H9 WT, H9-smDEL, H9-lgDEL, CT2 WT, CT2-smDEL, and CT2-lgDEL) were further engineered to contain a stably integrated cassette allowing for rapid induction of neurons using human NGN2 (Fernandopulle et al., 2018)(Methods)(Figure 1A). Neurons generated using this approach did not have any noticeable phenotypic differences between any of the deletions and controls in either background (Supplemental Figure 2). Examination of RNA-Seq signals in neurons generated by the inducible neuron system at the PWS locus confirmed the size of each deletion and targeting of the paternal allele due to lack of expression from the deleted region (Figure 1B). Analysis of gene expression between neurons and wild type hESCs revealed largely the same differentially expressed genes (DEGs) (Figure 1C, Supplemental Figure 2). Gene ontology analysis of the shared upregulated DEGs in WT, smDEL, and lgDEL neurons showed enrichment of terms of neuron-related processes, components, and function (Figure 1D).

**Figure 1.**
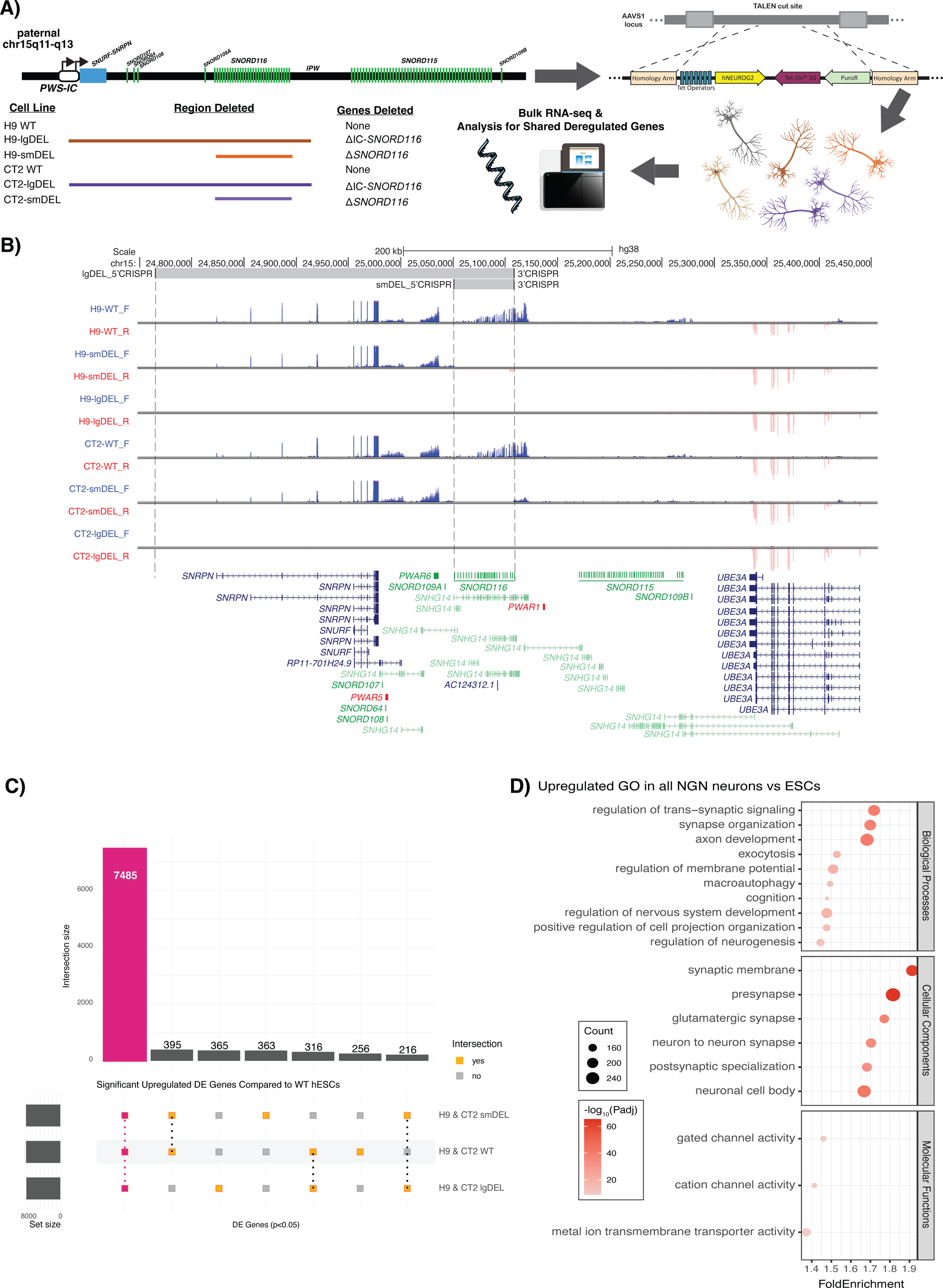
**A)** A schematic of our model system and experimental design. **B)** UCSC Browser image of the chromosome 15q11-q13 locus displaying representative bigwig tracks from each genetic background and genotype. Blue tracks show RNA signal from the sense (plus) strand; red tracks show RNA signal from the antisense (minus) strand. Top track shows CRISPR gRNA binding locations; gray shading indicates deleted region. GENCODEv25 gene annotations are shown at the bottom; protein-coding genes are shown in blue, noncoding genes are shown in green, and To Be Experimentally confirmed (TEC) biotype genes are shown in red. Some isoforms are removed for clarity. **C)** Upset plot comparing significant DEGs (p.adjust < 0.05) of wild type (WT), smDEL, and lgDEL inducible neurons across both genetic backgrounds to wild type H9 ESCs. Red bar represents significant shared upregulated (log_2_FoldChange > 0) DEGs in all three genotypes (WT, smDEL, and lgDEL) versus WT ESCs. **D)** Dot plot displaying gene ontology (GO) results for shared upregulated genes in all three genotypes versus WT ESCs. The x-axis represents the fold enrichment value, and y-axis shows ontology terms. Size of the dot corresponds to the number of DEGs in our data set contained within each ontology term. Shading of the dot corresponds to the negative log_10_ of the adjusted p-value, with more significant values shown in a darker shade.

### Eliminating expression from *SNHG14* promoters results in expression changes consistent with PWS phenotypes

Having confirmed that each of the lines harbored the desired deletions and generated neurons reliably, we set out to compare gene expression patterns across neurons. DEG analysis (Methods)(Supplemental Figure 3) of the lgDEL model neurons compared to WT neurons identified 483 upregulated DEGs and 381 downregulated DEGs shared across genetic backgrounds (Figure 2A)(Supplemental Table 2). This was a ∼5-fold and ∼3-fold enrichment of shared DEGs based on random permutations of similarly sized gene lists, respectively (*p* < 0.0001)(Supplemental Figure 3). When we inspected the PWS locus specifically, genes in the deletion were significantly differentially expressed as expected. However, we also noticed genes outside the boundaries of the engineered deletion were differentially expressed (Figure 2B, Supplemental Figure 3)(Supplemental Table 3). When we compared magnitudes of differential expression of all shared DEGs (483 upregulated, 381 downregulated) genes within and surrounding deletions of the PWS locus were most strongly affected (Figure 2C-D). Gene ontology analysis (Methods) on all 864 dysregulated genes revealed Molecular Function category terms related to ribosome structure, rRNA binding, and mRNA 5’-UTR binding, among others (Figure 2E, Supplemental Table 4). Interestingly, the DEGs present in the Structural Constituent of Ribosome category seem to be enriched for genes with lower expression in the brain compared to other tissue types (Supplemental Figure 4)(Rouillard et al., 2016). While disease ontology analysis on the 483 shared upregulated DEGs only returned two significant terms (Supplemental Figure 4), analysis of the 381 shared downregulated DEGs resulted in ontology terms related to phenotypes seen in PWS patients, such as delayed puberty, abnormality of the genital system, and obesity (Figure 2F, Supplemental Table 5). These results support the relevance of the lgDEL model in studying PWS.

**Figure 2.**
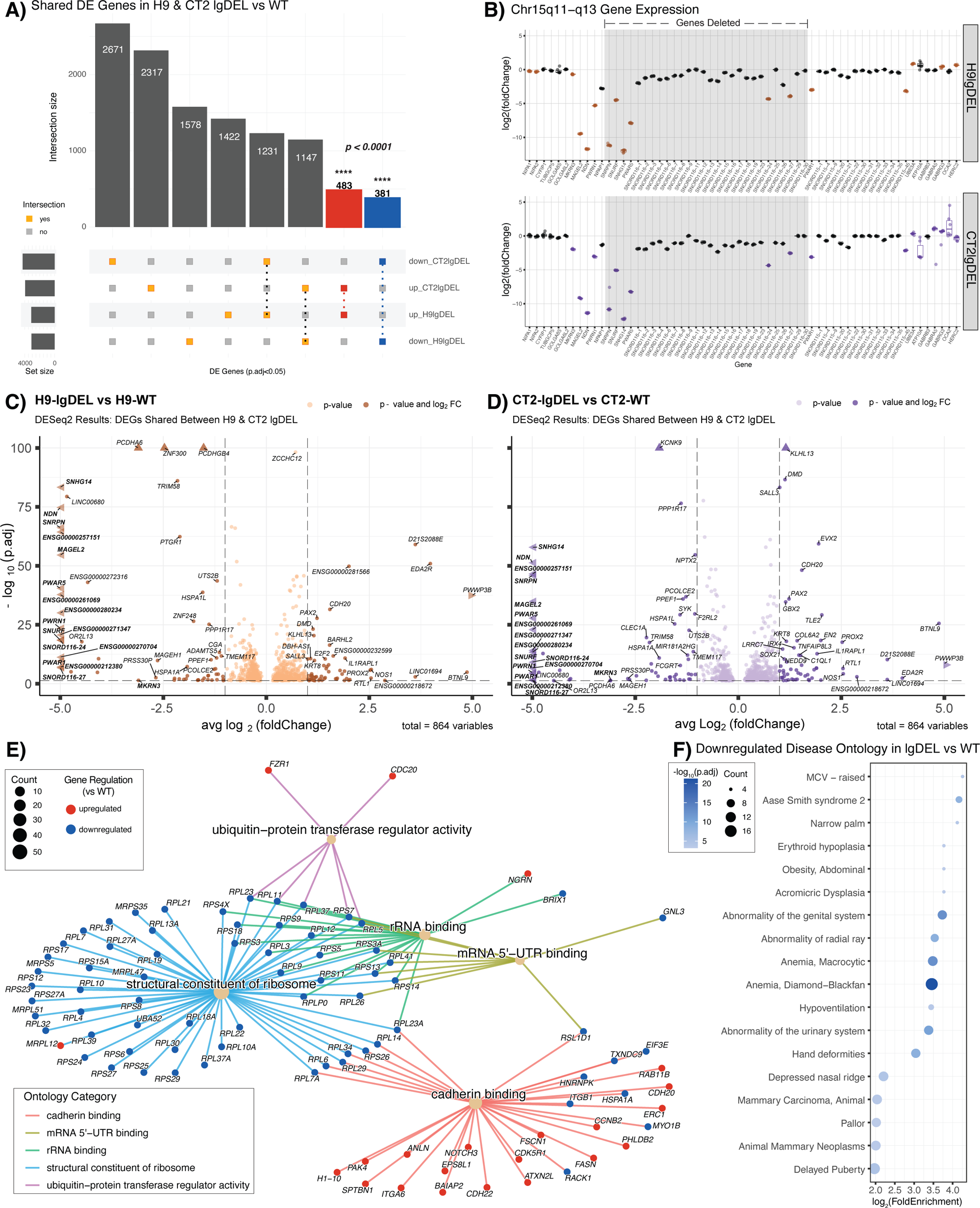
**A)** Upset plot comparing significant DEGs (p.adjust < 0.05) for lgDEL lines in both genetic backgrounds versus their isogenic WT controls. Red bar represents significant shared upregulated (log_2_FoldChange > 0) DEGs; blue bar represents significant shared downregulated (log_2_FoldChange < 0) DEGs. Significance of overlaps (p < 0.0001) determined via a permutation test. **B)** Box and whisker plots showing differential expression of genes of interest in chromosome 15q11-q13 region. Pseudocount was added to counts of all genes prior to calculation of log_2_(foldChange). Significant DEGs (p.adjust < 0.05) are shown in color, orange (H9lgDEL) or purple (CT2lgDEL). Gray shading indicates deleted region. **C&D)** Volcano plots displaying 864 significant (p.adjust < 0.05) DEGs shared between H9 & CT2 lgDEL lines. The x-axis represents the average log2 fold change of H9 & CT2 lgDEL lines, and y-axis represents the negative log_10_ of the adjusted p-value for C) H9 or D) CT2. Points in C) dark orange or D) dark purple correspond to DEGs with an average log_2_ fold change of < −1 or > 1. Genes contained within the chromosome 15q11-q13 region are denoted in bold. Points denoted by a triangle extend past the margins of the plot. **E)** Gene-concept network plot displaying GO terms of the molecular function (MF) category for all shared dysregulated genes. Main nodes (tan) correspond to the MF category with colored lines connecting to nodes of genes found in each category. Size of the main node corresponds to the number of DEGs in our data set contained within each ontology term. Colors of the gene nodes correspond to the log_2_ fold change for each DEG; red indicates log_2_FoldChange > 0, blue indicates log_2_FoldChange < 0. **F)** Dot plot displaying disease ontology results for shared downregulated genes. The x-axis represents the log_2_ fold enrichment value, and y-axis shows disease ontology terms. Size of the dot corresponds to the number of DEGs in our data set contained within each ontology term. Shading of the dot corresponds to the negative log_10_ of the adjusted p-value, with more significant values shown in a darker shade.

### Deletion of *SNORD116* alone is necessary to determine the targets and functions of *SNORD116* snoRNAs

While a large deletion model is relevant to many PWS cases, a recent report of a microdeletion encompassing the *SNORD116* cluster suggests these genes may be the primary contributor to the PWS phenotype (Tan et al., 2020). Therefore, we made a targeted deletion of the *SNORD116* C/D box snoRNA cluster (smDEL) that retains expression of the *SNHG14* parent transcript and *SNURF*-*SNRPN*. DEG analysis performed in a similar fashion as above (Methods)(Supplemental Figure 5)(Supplemental Table 6) revealed 178 upregulated DEGs and 139 downregulated DEGs shared across genetic backgrounds of our smDEL models (Figure 3A), a ∼7-fold and ∼9-fold enrichment of shared DEGs versus random permutations respectively (*p* < 0.0001)(Supplemental Figure 5). Similarly to the lgDEL model, the smDEL also impacted gene expression in the PWS locus beyond the bounds of the deletion (Figure 3B, Supplemental Figure 5)(Supplemental Table 7) and these were some of the most strongly effected genes (Figure 3C-D). However, the reduced number of genes resulted in fewer relevant gene ontology categories (Supplemental Table 8). Surprisingly, this reduced set was enriched for disease ontology terms related to phenotypes seen in PWS patients, like short toe and short palm (Figure 3D, Supplemental Table 9)(Cassidy & Driscoll, 2009).

**Figure 3.**
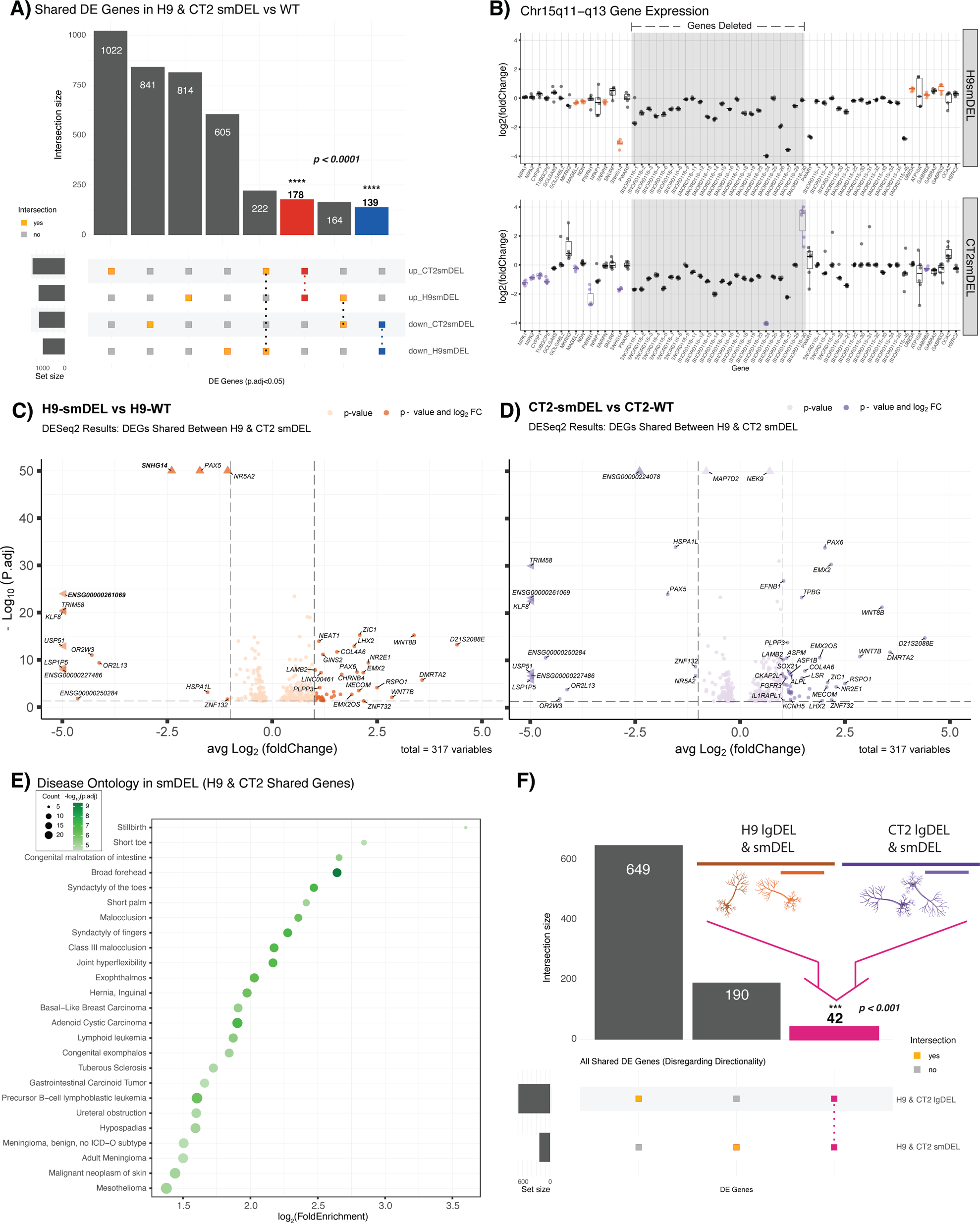
**A)** Upset plot comparing significant DEGs (p.adjust < 0.05) for smDEL lines in both genetic backgrounds versus their isogenic WT controls. Red bar represents significant shared upregulated (log_2_FoldChange > 0) DEGs; blue bar represents significant shared downregulated (log_2_FoldChange < 0) DEGs. Significance of overlaps (p < 0.0001) determined via a permutation test. **B)** Box and whisker plots showing differential expression of genes of interest in chromosome 15q11-q13 region. Pseudocount was added to counts of all genes prior to calculation of log_2_(foldchange). Significant DEGs (p.adjust < 0.05) are shown in color, orange (H9smDEL) or purple (CT2smDEL). Gray shading indicates deleted region. **C&D)** Volcano plots displaying 317 significant (p.adjust < 0.05) DEGs shared between H9 & CT2 smDEL lines. The x-axis represents the average log2 fold change of H9 & CT2 smDEL lines, and y-axis represents the negative log_10_ of the adjusted p-value for C) H9 or D) CT2. Points in C) dark orange or D) dark purple correspond to DEGs with an average log_2_ fold change of < −1 or > 1. Genes contained within the chromosome 15q11-q13 region are denoted in bold. Points denoted by a triangle extend past the margins of the plot. **E)** Dot plot displaying disease ontology results for all shared dysregulated genes. The x-axis represents the log_2_ fold enrichment value, and y-axis shows top 25 disease ontology terms. Size of the dot corresponds to the number of DEGs in our data set contained within each ontology term. Shading of the dot corresponds to the negative log_10_ of the adjusted p-value, with more significant values shown in a darker shade. **F)** Upset plot comparing all significant DEGs (p.adjust < 0.05) for both H9-smDEL & CT2-smDEL and H9-lgDEL & CT2-lgDEL versus isogenic WT controls. Pink bar represents significant DEGs dysregulated across both genetic backgrounds and deletion types versus isogenic WT controls. Significance of overlap (p < 0.001) determined via a permutation test.

### Comparison of small and large deletion models reveals a novel and robust regulatory network of genes consistently dysregulated in PWS-like systems

Having demonstrated that DEGs in each set of models identified genes enriched for PWS relevant phenotypes, we wondered if any DEGs – besides those within the PWS locus – might be shared across lgDEL and smDEL models. We hypothesized that genes shared across all comparisons are central to the disorder and therefore important to focus on. We further filtered our DEGs from the lgDEL and smDEL models (Methods)(Supplemental Figure 6)(Supplemental Tables 10-13) which resulted in 649 total DEGs in the lgDEL model and 190 total DEGs in the smDEL model. After overlapping these two lists, we found 42 genes shared between both genetic backgrounds and deletions (Figure 3F), a ∼3-fold enrichment of shared DEGs versus random permutations (*p* < 0.001)(Supplemental Figure 6). The list of 42 genes contains 8 transcription factors and 3 genes located within the PWS locus at chr15q11-q13 (Supplemental Table 14). While binding profiles of these transcription factors have not been studied in the context of PWS, we turned to the Enrichr gene set enrichment database that has compiled many different resources of experimental and predicted DNA binding and protein-protein interactions (Chen et al., 2013; Kuleshov et al., 2016). Specifically, we queried the Enrichr_Submissions_TF-Gene_Cooccurrence library, which has been compiled from over 300,000 gene set submissions, to evaluate the co-occurrence of our shared genes and transcription factors. This approach has proven effective in both identifying established gene interactions and uncovering new ones (Ma’ayan & Clark, 2016). When we analyzed the set of 42 shared genes, we found that 6 out of the 8 TFs in the shared gene list showed significant co-occurrence (Supplemental Table 15). Further, disease ontology analysis on the 42 shared genes (Methods) revealed among the most significant ontology categories were those associated with Intellectual Disability/Mental Retardation (Figure 4A)(Supplemental Table 16), a trait commonly associated with PWS (Cassidy & Driscoll, 2009). Though we analyzed gene expression in a neuronal model, many of the disease ontology enrichments we obtained are not directly related to neuronal function. When we examined expression of the 42 shared genes across dozens of tissues profiled by the Genotype-Tissue Expression (GTEx) project (https://gtexportal.org/home/multiGeneQueryPage)(Carithers et al., 2015), we noticed many of these genes were expressed across multiple tissue types, not just the brain, suggesting they might be co-expressed in different contexts and may physically interact (Supplemental Figure 7). To determine potential interactions between the resultant protein products of the 42 shared genes, we utilized the STRING database (v.11.5, https://string-db.org/)(Szklarczyk et al., 2015). We found that our shared gene network had 24 more edges than the expected value with a PPI enrichment p-value of 0.00276, which means our protein network was predicted to have more interactions than expected for a random set of the same size and distribution across the genome (Supplemental Figure 7). In addition, these genes had significantly lower median LOEUF score, a measure of a gene’s likelihood to have a deleterious mutation in the healthy population, compared to the remainder of the genes contained within the gnomAD database (v.2.1.1, https://gnomad.broadinstitute.org/)(Karczewski et al., 2020)(Supplemental Figure 7), further supporting the potential disease relevance of this gene network.

**Figure 4.**
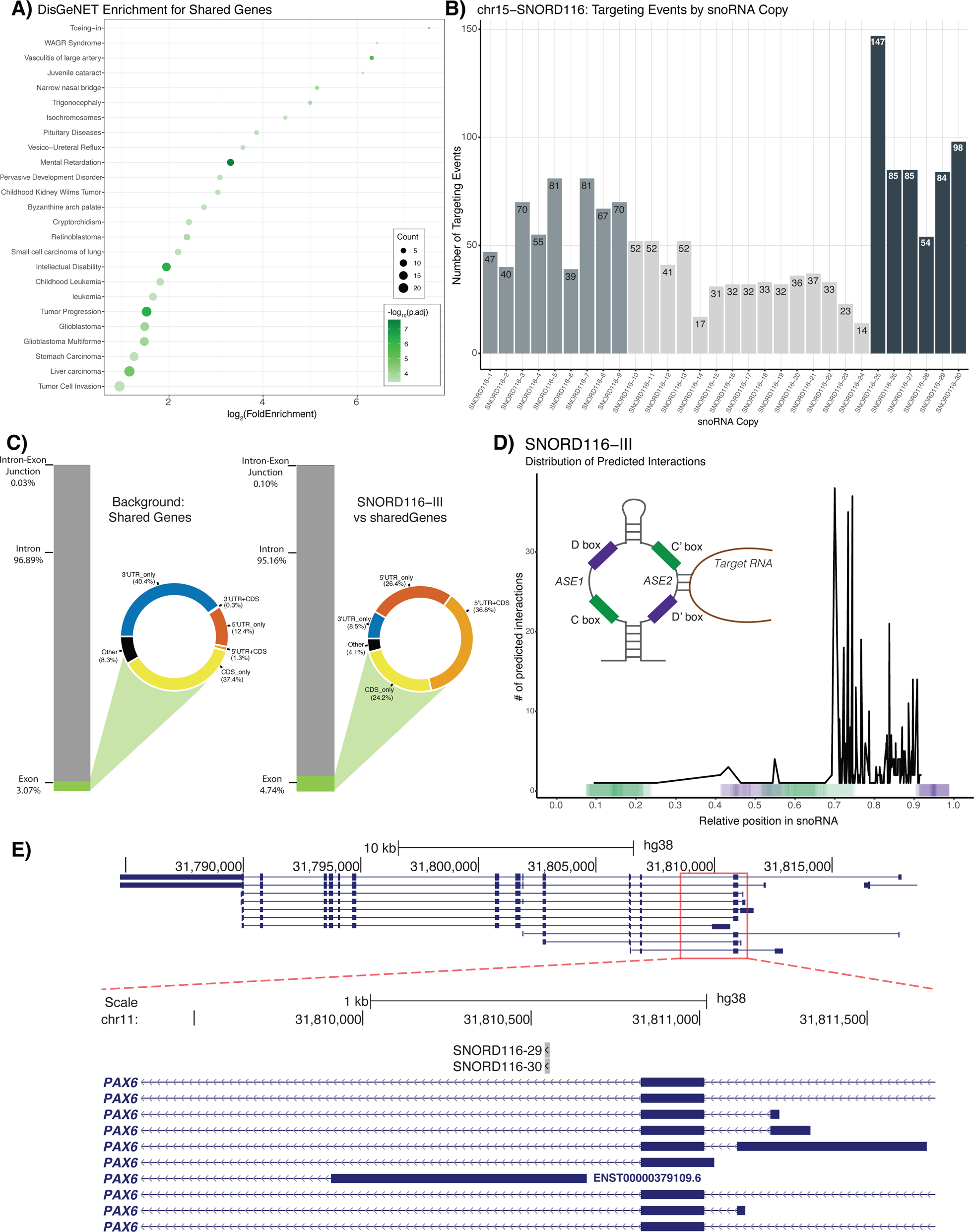
**A)** Dot plot displaying DisGeNET results for 42 shared dysregulated genes. The x-axis represents the log_2_ fold enrichment value, and y-axis shows top 25 ontology terms. Size of the dot corresponds to the number of DEGs in our data set contained within each ontology term. Shading of the dot corresponds to the negative log_10_ of the adjusted p-value, with more significant values shown in a darker shade. **B)** Bar plot represents the number of predicted targeting events per copy of SNORD116. Colors of the bars correspond to the three subgroups of *SNORD116*: *SNORD116-I* (*SNOG1*, copies 1-9), *SNORD116-II* (*SNOG2*, copies 10-24) and *SNORD116-III* (*SNOG3*, copies 25-30). **C)** Bar charts representing the nucleotide composition of the shared 42 dysregulated genes (Background: Shared Genes) and the nucleotide composition of the predicted targeting events by *SNORD116-III* copies on those shared genes (*SNORD116-III* vs Shared Genes) by genomic feature category: exon, intron, and intron-exon junctions. Exon category is subdivided based on genic location and displayed as a donut plot. Coloring of donut plots is based on exon category; 5’UTRs are represented in orange, 3’UTRs are represented in blue, CDS is represented in yellow, and any portion of exonic sequence not falling under those categories is termed “other” and shown in black. **D)** Plot displaying distribution of prediction interactions for *SNORD116-III*. The x-axis corresponds to the relative position within snoRNA copies, and y-axis represents the number of predicted interactions for which the center of the predicted binding interaction was used (black line). Color-coded bar on the x-axis indicates the position of C/C’ and D/D’ boxes found in snoRNA copies, indicated by green and purple respectively. **E)** UCSC Browser image of the *PAX6* locus displaying BEDtracks of *SNORD116-III* predicted binding. Top track shows entire *PAX6* gene. Bottom track shows zoomed in view, with predicted binding shown to occur in 5’UTR of one transcript of *PAX6* (ENST00000379109.6).

### *SNORD116* snoRNAs are predicted to directly regulate a subset of our novel gene network

Given results above suggested that these genes interact with each other in multiple ways, including transcriptional regulation and protein-protein interactions, we wondered whether these genes may be directly regulated by *SNORD116* snoRNAs. We employed a novel C/D box snoRNA prediction tool, snoGloBe (Deschamps-Francoeur et al., 2022), which predicted a significant enrichment of *SNORD116* interactions with our shared gene list versus several control analyses. Examining the distribution of these predicted targeting events revealed that 35 of the 42 genes are predicted to be targeted by *SNORD116* (Supplemental Figure 8)(Supplemental Table 17). When we plotted the number of predicted binding events per copy of *SNORD116*, we observed a correlation between the number of predicted binding events and the established breakdown of *SNORD116* snoRNAs into its three subgroups: Group I (*SNOG1*, *SNORD116-1* to *SNORD116-9*), Group II (*SNOG2*, *SNORD116-10* to *SNORD116-24*), and Group III (*SNOG3*, *SNORD116-25* to *SNORD116-30*)(Castle et al., 2010; Runte et al., 2001)(Figure 4B). Interestingly, we noted that *SNORD116* Group III copies, referred to henceforth as *SNORD116-III,* showed the highest number of predicted binding events per copy.

To analyze the significance of our results, we first compared the number of predicted targeting events per snoRNA copy of *SNORD116* to *SNORD115* (Supplemental Figure 8)(Supplemental Table 18) and saw that *SNORD116* copies have an enrichment of predicted targeting events per copy. Additionally, genes with predicted targeting events were significantly enriched for predicted targeting by *SNORD116-III* versus *SNORD115* copies (Supplemental Figure 9). Compared to a random permutation (Methods), we observed a significant ∼2.5-fold enrichment of the mean, median, and sum of *SNORD116-III* predicted targeting events on the shared gene list (*p* < 0.01)(Supplemental Figure 10). Similarly to the findings presented by Deschamps-Francoeur et al., when we compared the background genomic feature coverage of our shared genes list to the genomic feature coverage of *SNORD116-III* predicted binding events, we saw an enrichment in both exon and intron-exon junction categories. Most notably, there was a large increase in coverage of 5’-UTRs (Figure 3D)(Supplemental Figure 11), which may suggest a role for *SNORD116* in regulation of translation of the shared dysregulated genes. Finally, when we examined the distribution of the predicted binding events across snoRNA copies (Methods)(Figure 4D), we obseved predicted binding events for *SNORD116-III* copies mainly occur upstream of the D’ box at the second antisense element (ASE2), a portion of this class of snoRNA that typically interacts with target RNAs (Kiss-László et al., 1996; Nicoloso et al., 1996). This trend is less clear for other *SNORD116* groups and for our control *SNORD115* copies, which show a greater portion of predicted targeting events occuring in the C/C’ boxes (Supplemental Figure 12). An example of *SNORD116-III* predicted binding within a 5’-UTR of one of our shared DEGs, *PAX6*, is shown in Figure 4E. This 5’-UTR is annotated in transcript ENST00000379109, an alternative form of the canonical *PAX6* transcript, which contains 422 amino acids.

## Discussion

While it has long been understood that perturbations of the chr15q11-13 region cause PWS, it is unclear if the genes included in the deletions are directly related to PWS phenotypes, if genes regulated by them are to blame, or if it is some combination of these effects. Multiple studies have attempted to address this issue by characterizing gene expression in postmortem PWS brain tissues and neurons differentiated from PWS patient-derived pluripotent stem cell lines to identify genes dysregulated in this disorder (Bochukova et al., 2018; Falaleeva et al., 2015; Huang et al., 2021; Sledziowska et al., 2023; Victor et al., 2021). While these studies indicate gene expression is indeed dysregulated in PWS patient samples, our analysis here showed few genes had consistent dysregulation across a subset of these studies (Supplemental Figure 1A). Furthermore, the genes that showed consistent trends across these studies seemed to have limited relevance to PWS based on gene ontologies (Supplemental Figure 1B). This discordance in gene expression patterns could be attributed to multiple reasons, both technical and biological. Obtaining controls from otherwise healthy donors for postmortem brain tissue comparisons matched for age, sex, genetic background, and postmortem delay is extremely challenging. For iPSC-based, experiments the background of genetic variants outside of the chr15q11-13 region could be substantially different between PWS patients and otherwise healthy controls. This is problematic as multiple studies have established that genetic background of induced pluripotent stem cells (iPSCs) can contribute substantially to changes in gene expression (Banovich et al., 2018; DeBoever et al., 2017; Kilpinen et al., 2017; Rouhani et al., 2014; Thomas et al., 2015). Even heterogeneity found in neuronal differentiation of these cellular models can prove to be a challenge in generating reproducible differential gene expression results (Zeng et al., 2010). These background effects could be potentially mitigated in PWS patient derived cells if the missing genetic material could be restored. However, the size of the deletions frequently present in PWS patients poses a challenge for replacing missing genetic information to generate such isogenic controls.

To combat these issues, we utilized multiple isogenic cell lines and an inducible differentiation protocol to generate reproducible, homogenous neurons. The caveats of this system include a lack of electrically active neurons, a more artificial path through neuronal differentiation, and that these are cortical neurons, as opposed to hypothalamic neurons which are most often implicated in PWS physiology (reviewed in Swaab, 1997). The lgDEL model harbors a deletion encompassing all promoters of the *SNRPN* transcript, which eliminates transcription of the host gene and therefore processing and expression of *SNORD116*. The smDEL model harbors a targeted deletion of just the *SNORD116* snoRNAs, designed to model the smallest known deletion to still result in PWS phenotypes (Tan et al., 2020). As *SNORD116* snoRNAs are not polyadenylated and thus not enriched for during polyA-RNA-Seq, most *SNORD116* copies do not meet cutoffs to be called DEGs in our data set (Figure 2B, 3B). However, the lack of signal from the *SNORD116* locus demonstrated successful deletion of the region in both models (Figure 1D).

Notably, in the lgDEL model we saw differential expression of a subset of ribosomal protein genes. While these genes are typically thought to be utilized similarly across most tissues, the sets of ribosomal DEGs identified here have generally lower expression in brain compared to other tissues profiled by GTEx (Supplemental Figure 4A). This could suggest that due to their lower starting expression levels, these proteins are more sensitive to small perturbations. As neither *SNURF* nor *SNRPN* are significantly dysregulated in the smDEL model, we believe this analysis may demonstrate separable functions of *SNURF-SNRPN* and *SNORD116* snoRNAs. Specifically, *SNURF* and/or *SNRPN* may have a specialized role in ribosomal gene expression while the *SNORD116* snoRNAs may have a completely different role. C/D box snoRNAs have generally been shown to bind and modify ribosomal RNAs (Decatur & Fournier, 2002; T. Kiss, 2001). However, both *SNORD116* and *SNORD115* snoRNA gene clusters in the chr15q11-13 region are known as orphan snoRNAs and do not show any sequence homology with rRNAs. Previous studies have predicted binding events of these snoRNAs using basic sequence matching approaches, however these results have not been confirmed in a disease-relevant context (Baldini et al., 2022; Bazeley et al., 2008; Kehr et al., 2011). Upon further investigation, none of the genes previously predicted to be targeted by *SNORD116* (Baldini et al., 2022) were differentially expressed in our smDEL model across both genetic backgrounds. More recent snoRNA prediction tools have employed machine learning techniques trained on large scale RNA-RNA interaction data to develop models for systematic prediction of such interactions (Deschamps-Francoeur et al., 2022). Application of this tool to the consistently differentially expressed genes revealed significantly elevated numbers of predicted targeting events by *SNORD116*, particularly amongst group III copies. Importantly we leveraged *SNORD115* copies as controls in this analysis. As *SNORD115* is also a cluster of C/D box snoRNAs contained within the same locus and its deletion alone has no observable phenotypes (Runte et al., 2005), it serves as a relevant comparator. The predicted *SNORD116* binding sites were facilitated primarily by the second antisense element of *SNORD116* sequences, consistent with described mechanisms of C/D box snoRNA targeting (Kiss-László et al., 1996; Nicoloso et al., 1996). The predicted binding sites on the consistently dysregulated genes were particularly enriched at 5’-UTR regions suggesting a potential role in modulating transcript stability and/or translation (Hinnebusch et al., 2016). Even though we observed a slight enrichment of predicted binding at intron-exon junctions and snoRNAs have been implicated in alternative splicing (Falaleeva et al., 2016; Scott et al., 2012), we do not believe this small enrichment suggests a significant role for *SNORD116* in splicing. Additionally, our analysis suggests that even amongst *SNORD116* there is bias in gene regulation (Figure 4B). The group III copies have been reported to be absent from the rodent lineage (Baldini et al., 2022) and could begin to explain differences in phenotypes observed in mouse models of PWS. Subsequent targeted deletions of individual *SNORD116* groups could shed more light on these findings.

The novel list of genes we have described holds promise for future studies. There are a number of fascinating transcription factors, like *PAX6* which may contribute to some of the vision phenotypes reported in PWS patients (Bohonowych et al., 2021); *IRX5* which has been implicated in obesity and metabolism (Sobreira et al., 2021); and *FGF13* which has links to developmental delay (Fry et al., 2021). Most notably, however, is the consistent dysregulation of *MAGEL2*. Mutations in *MAGEL2* cause Schaaf-Yang syndrome (SYS), which shares some phenotypes with PWS. Even more interesting is that *MAGEL2* is the only shared gene across the subset of previously mentioned studies we analyzed and this study. This may suggest that both *SNORD116* loss and *MAGEL2* dysregulation drive PWS phenotypes. While we have endeavored to create a well-controlled experimental design at the genetic level, a significant limitation of this study is that we are unable to differentiate between the effects of loss of *SNORD116* expression and loss of the genetic region itself. As mentioned above, the *SNORD116* DNA sequences may play a role in silencing of the locus. Furthermore, other work from our group indicated regions such as *IPW* can form long range interactions to *MAGEL2* and other surrounding genes (Hsiao et al., 2019). Thus, the deletions we have constructed, even the smallest one, could have large scale impacts on chromatin organization and merit further investigation.

## Materials and Methods

### Genome editing of hESCs

H9 ESCs were first engineered with a deletion of the entire *SNHG14* transcript (lgDEL) or *SNORD116* alone (smDEL) and then subsequently edited to introduce a neurogenin-2 (NGN2) cassette into the AAVS1 locus following the protocol described below.

#### Preparation

Guide RNAs for the lgDEL and smDEL were designed using available guide RNA design tools (Supplemental Table 1). Each guide was cloned into the pSpCas9(BB)-2A-Puro (PX459) V2.0 plasmid, a gift from Feng Zhang (Addgene, #62988). This plasmid was digested with Bbs1 restriction enzyme and ligated with the guide RNA insert.

Two days prior to planned genome editing, a 100mm dish of mitotically inactivated DR4 mouse embryonic fibroblasts (MEFs) was prepared. hESCs were gown on mitotically inactivated MEFs and fed daily with sterile-filtered DMEM/F12 media (Gibco, # 11330032) supplemented with 20% Knock Out Serum Replacement (Gibco, #10828028), 1X MEM Non-essential amino acids (Gibco, #11140050), 1mM L-glutamine (Gibco, #25030081) with 0.14% β-mercaptoethanol, and bFGF (Gibco, #PHG0023), until ∼60-75% confluent. Cells were treated with 10uM ROCK inhibitor, Y-27632 2HCl (Tocris #1254), 24 hours prior to planned genome editing.

#### Nucleofection

The day of editing, approximately 1-1.5 × 10^6^ cells were treated with Accutase (Millipore, SCR005) to release the cells from the plate, cell suspension was singularized by pipetting and then pelleted. The media was removed from the cell pellet and cells were resuspended according to the protocol provided for the P3 Primary Cell 4D-Nucleofector Kit (Lonza, V4XP-3024). Briefly, a mixture of 82uL nucleofector solution, 18uL nucleofection supplement, and desired plasmids were added to the pellet. The pellet was resuspended in the solution by pipetting gently three times using a P200 pipet. For the smDEL and lgDEL edits, 2.5ug of each CRISPR plasmid was added to the nucleofection solution (Supplemental Table 1). For the introduction of the NGN2 cassette, 2μg of both TALEN-L and TALEN-R plasmids (Addgene, #59025 and #59026) and 4µg of pUCM-AAVS1-TO-hNGN2 plasmid (Addgene, #105840) was added to the nucleofection solution. The cell suspension was transferred to the nucleofection cuvette using the P200 pipet and nucleofection was performed on the 4D-Nucleofector (Lonza) on the program for hESC, P3 primary cell protocol. After nucleofection, hESC suspension was transferred to the 100mm dish plated with DR4 MEFs containing the KOSR media mentioned above supplemented additionally with 10uM ROCK inhibitor using the transfer pipet included in the Lonza kit. For the NGN2 edit, the media of the 100mm dish was also supplemented with 5µM L755507 (Selleckchem, #S7974) and 1µM SCR7 (Selleckchem, #S7742) to encourage homology directed repair to incorporate the NGN2 cassette into the AAVS1 locus.

#### Selection

For lgDEL and smDEL edits, feeding media was changed 24 hours following transfection (Day 1 post-transfection) and supplemented with 1 ng/μL puromycin and 10μM ROCK inhibitor. This selection was continued for 48 hours total to select cells transiently expressing the vectors containing the gRNA and Cas9 protein. On Day 2, the media was changed and supplemented with fresh 1μg/μL puromycin and ROCK inhibitor. On Day 3, the media was changed and supplemented with fresh ROCK inhibitor.

Subsequent media changes occurred every other day, supplemented with fresh ROCK inhibitor. Once small colonies became visible, media changes occurred daily with fresh media alone. After a total of 15 days, each colony was manually passaged into its own well of a 24-well plate coated with mitotically inactivated MEFs via cutting and pasting. Feeding media in the 24-well plate was supplemented with 10uM ROCK inhibitor to encourage cell attachment. 48 hours after passaging cells, the feeding media was changed. Approximately 4 days after passaging to a 24-well plate, a few colonies from each well were isolated into PCR tube strips and pelleted for screening.

For the NGN2 edit, feeding media was changed 24 hours following transfection supplemented with fresh 10µM ROCK inhibitor, 5µM L755507, and 1µM SCR7. Between 72-96 hours post-transfection, selection began by supplementing fresh feeding media with 1µg/μL puromycin and 10µM ROCK inhibitor. Selection continued for 4 or 5 days by changing feeding media and supplementing with fresh 1µg/μL puromycin. After selection, colonies were grown to a size sufficient for clonal isolation. Each colony was manually passaged into its own well of a 24-well plate coated with mitotically inactivated MEFs via cutting and pasting. After approximately one week of growth, a few colonies from each clone were manually passaged to a new 24-well plate. The remaining colonies from each clone were transferred to a 1.5-mL microcentrifuge tube and pelleted for screening.

#### Screening

For lgDEL and smDEL edits, DNA was extracted using the HotSHOT method (Truett et al., 2000). In brief, media was removed from pelleted cells and 30µL of alkaline lysis buffer (25mM NaOH, 0.2mM EDTA, pH ∼12) was added to each tube. The tubes were incubated at 95°C for 45 minutes. Subsequently, 30µL of neutralization reagent (40mM Tris-HCl, pH ∼5) was added to each tube. Tubes were capped tightly, flicked to mix, and spun down.

For the NGN2 edit, genomic DNA (gDNA) was extracted using a homemade lysis buffer containing 1% SDS, 75mM NaCl, 25mM EDTA, and 200μg/mL Proteinase K. Briefly, 500µL of the lysis buffer was added to each cell pellet and the tubes were incubated at 63°C overnight. The following day, 170μL of 150mM NaCl was added, followed by the addition of 670μL of chloroform. The mixture was shaken vigorously (∼60 times) and centrifuged at 11,000 rcf for 10 minutes at room temperature. The top aqueous layer (∼650µL) was removed and transferred to a new tube to which an equal amount of 100% isopropanol was added. The mixture was shaken ∼10 times and was incubated at −20°C for 20 minutes. Next, the mixture was centrifuged at 20,000 rcf for 20 minutes at 4°C. The supernatant was removed, and the pellet was washed with 70% ethanol before being resuspended in 50µL of 10mM sterile-filtered Tris.

Genotyping was performed using the Herculase II Fusion DNA Polymerases kit (Agilent, #600677) following manufacturer’s protocol. For the DNA template, 1µL of each sample was used per 25µL reaction. The annealing temperature was 60°C for all primer combinations (Supplemental Table 1). The PCR products were run on a 1% agarose gel for 35 min at 95V. Primer sets were designed upstream of the 5’ CRISPR cut site and downstream of the 3’ CRISPR cut site for each lgDEL and smDEL edits. For lgDEL and smDEL clone screening, first PCR primers for knockout of the region of interest were utilized (lgDEL or smDEL, primer set 1,2). If there was successful knockout on one or both alleles, a band would be present. Any clones identified as positive for a knockout then were screened using PCR primers to identify heterozygous clones (lgDEL and smDEL, primer set 1,2). For a heterozygous clone, a band would be present. RNA was extracted from heterozygous clones and subjected to cDNA synthesis using the SuperScript First-Strand Synthesis System for RT-PCR (Invitrogen, #11904018) following manufacturer’s protocol to test for the parent-of-origin of the deleted allele. Finally, RT-PCR was performed on the cDNA with primers for *SNORD116* and a control, *GAPDH* (Supplemental Table 1). Clones which did not express *SNORD116* were then further expanded and banked down. One such clone from each genotype was subsequently edited for incorporation of NGN2. For NGN2 edits, a nested PCR across the insertion sites was used to identify clones which NGN2 was incorporated into the AAVS1 locus in the correct orientation. These primers were designed so that one primer was in the endogenous AAVS1 locus and the other primer was in the exogenous transgene. Following the first PCR (PCR1, using primer sets 1,2 and 3,4), a second “nested” PCR (PCR2, using primer sets 1,2 and 3,4) was run utilizing the product from PCR1 as the template for PCR2. Final nested products from both primer sets with banding at ∼1 kb indicated successful incorporation of the NGN2 cassette. Clones with the NGN2 cassette integration were further screened for heterozygosity utilizing primers for the wild type AAVS1 locus (primer set 1,2). Wild type or heterozygous clones would show a band at 500 bp. Clones which showed correct, homozygous insertion of NGN2 were expanded and banked down. One such clone from each genotype was subsequently used for the sequencing experiments described.

### hESC culture

To transition cells from feeder conditions to feeder-free conditions, cells were manually passaged by cutting and pasting colonies once confluent. After 5-7 days, any differentiation was manually removed before first passage. Routine culture of H9 and CT2 ESCs was done using feeder-free conditions. Cells were maintained in mTeSR™ Plus media (STEMCELL Technologies, Catalog #100-0276) on Matrigel™ hESC-Qualified Matrix (Corning™, Catalog #354277) coated 6-well plates in a humidified atmosphere with 5% CO_2_ at 37°C. Feeding media was changed daily. Cells were passaged once 80-100% confluency was reached, approximately every 4-5 days. Briefly, media was removed from well(s), well(s) were gently rinsed with sterile PBS, sterile filtered 0.5 mM EDTA in PBS was added to well(s), and the plate was placed back into the incubator undisturbed for 2-5.5 minutes. After incubation, EDTA solution was gently aspirated from well(s), being careful to not disturb cells. Using a 2-mL serological pipette, 1 mL of media was added to well(s) while gently scraping bottom of well(s) to dislodge cells. The cell suspension was pipetted 1-2 times to break up the cells into clumps. 75-125 µL of cell suspension was added to a new well containing 2 mL of culture media supplemented with 10uM ROCK inhibitor.

### Inducible neuron differentiation

hESCs were differentiated into cortical neurons following an established protocol (Fernandopulle et al., 2018) with some modifications. When hESCs reached 70-80% confluency, cells were prepared for differentiation. First, any differentiated cells were manually removed, and wells were gently rinsed with sterile PBS. Cells were treated with Accutase and the plate was placed in the incubator for 2 minutes. The plate was agitated as needed during the incubation time encourage release of the cells from the plate. After incubating, 1 mL of media was added to cell suspension and cells were singularized by pipetting with a 2-mL serological pipet. Cell suspension was transferred to a 15-mL conical tube and centrifuged at 1,200 rpm for 3 minutes. Media was aspirated from pellet and pellet was resuspended in Induction Media (IM). IM was prepared by supplementing DMEM/F12 with HEPES (Gibco, #11330032) with 1X N2 supplement (Gibco, # 17502048), 1X MEM Non-essential amino acids, and 1X GlutaMAX (Gibco, #35050061). Cells were counted using a hemocytometer and plated for differentiation in IM supplemented with 10µM ROCK inhibitor and 2µM doxycycline hydrochloride (Fisher Scientific, BP2653-5). Cells were fed daily with IM supplemented with 2uM doxycycline hydrochloride for 3 days. On day 4 of differentiation, cells were again singularized with Accutase as above. The cell pellet was resuspended in Cortical Media (CM) supplemented with 10µM ROCK inhibitor. CM was prepared by mixing equal amounts of DMEM/F12 with HEPES and Neurobasal Medium (Gibco, #21103049) and adding 1X B27 supplement (Gibco, #17504044), 10 ng/mL BDNF (R&D Systems, 248-BD), 10 ng/mL GDNF (R&D Systems, 212-GD), 10 ng/mL NT3 (PeproTech, 450-03), and 1 µg/mL laminin (Gibco, #23017015). Cells were counted using a hemocytometer and plated at 1 million cells per well of a 6-well plate or 7 million cells per 10-cm dish in CM supplemented with 10uM ROCK inhibitor. Plates and dishes were coated prior to plating with 100 µg/mL poly-D-lysine hydrobromide (Millipore, P0899) and 5 µg/mL laminin (Gibco, #23017015). A complete media change with CM was performed the following day. Media was changed every other day until collection on day 11.

### Cell collection

For hESCs, any differentiated cells were manually removed, and wells were gently rinsed twice with sterile PBS. Sterile filtered 0.5 mM EDTA in PBS was added to wells, and the plate was placed back into the incubator undisturbed for 5.5 minutes. After incubation, EDTA solution was gently aspirated from wells, being careful to not disturb cells. Using a 2-mL serological pipette, 1 mL of sterile PBS was pipetted down the back of wells to dislodge cells. The cell solution was transferred to a 1.5-mL microcentrifuge tube and centrifuged at 2000xg for 5 minutes at 4°C. PBS was aspirated from pellets. Pellets were flash frozen in liquid nitrogen and stored at −80°C until RNA extraction.

For day 11 hESC-derived neurons, media was aspirated from the wells/dish. DMEM/F12 was added to the wells/dish and the cells were scraped to detach them from the plate. The cell suspension was collected in a 15-mL conical tube and spun down at 2000 rpm for 3 minutes. Media was aspirated from pellet and pellet was resuspended in 1 mL of TRIzol™ (Invitrogen™, #15596026). The cell suspension was transferred to a 1.5-mL microcentrifuge tube. The tube was briefly vortexed and incubated at room temperature for 5 minutes before proceeding with RNA extraction.

### RNA extraction

For hESCs, RNA was harvested using the miRNeasy® Mini Kit (QIAGEN, #1038703) following manufacturer’s protocol with minor modifications. The work surface, pipettes, and centrifuge rotors were treated with RNAse Away (Life Technologies, #10328011) prior to beginning extraction. Pellets were transferred from storage at −80°C to ice. Samples were homogenized in 700µL QIAzol by pipetting and brief vortexing. Cell lysate was applied to QIAshredder columns (QIAGEN, #1011711). Samples were incubated at room temperature for 5 minutes. Following incubation, 140 µL of chloroform was added to the homogenate and shaken vigorously for 15 seconds. Samples were incubated at room temperature for 2-3 minutes and then centrifuged for 15 minutes at 12,000xg at 4°C. Approximately 400 µL of the aqueous phase was transferred to a new 1.5-mL microcentrifuge tube. A second chloroform extraction was performed by adding an equal volume of chloroform to the aqueous phase and shaking vigorously for 15 seconds. The samples were centrifuged for another 15 minutes at 12,000xg at 4°C, and the aqueous phase (∼350 µL) was transferred to a new 1.5-mL microcentrifuge tube to which 1.5 volumes of 100% ethanol was added. The contents of the tube were mixed by pipetting and applied to the RNeasy spin column, following manufacturer’s instructions for on-column DNase treatment using RNase-Free DNase Set (QIAGEN, #79254) and the addition of a second wash with Buffer RPE. For hESC-derived neurons, RNA was harvested using the Direct-zol RNA Miniprep kit (Zymo Research, Cat No. R2050) following manufacturer’s protocols. For both hESCs and hESC-derived neurons, RNA was eluted in RNase-Free water and stored at −80°C until library construction, for which 1 µg of RNA was used.

### RNA-seq library preparation and sequencing

Total RNA quality for hESC samples and most hESC-derived neuron samples was assessed using the Agilent TapeStation 4200 with RNA ScreenTape Analysis, including RNA ScreenTape (Agilent, 5067-5576), RNA ScreenTape Sample Buffer (Agilent, 5067-5577), and RNA ScreenTape Ladder (Agilent, 5067-5578). All samples measured had an RNA Integrity Number (RIN) of 8.4 or greater.

For hESCs, RNA libraries for RNA-seq were prepared using the NEBNext® Ultra II Directional RNA Library Prep Kit for Illumina® (NEB, #E7760L) following manufacturer’s protocol for use with NEBNext Poly(A) mRNA Magnetic Isolation Module (NEB, #E7490). Libraries were checked for quality and average fragment size using ScreenTape analysis, including D1000 ScreenTape (Agilent, 5067-5582) and D1000 Sample Buffer (Agilent, 5067-5602). Concentration of libraries was measured using Qubit™ 2.0 Fluorometer with Qubit™ dsDNA HS Assay Kit (Invitrogen™, Q32851). Molar concentration of libraries was determined using NEBNext® Library Quant Kit for Illumina® (NEB, #E7630) following manufacturer’s protocol for 6 Standards, 100–0.001 pM (Section 3). Quantification of libraries was calculated using the worksheet from NEBioCalculator (v1.15.0, https://nebiocalculator.neb.com/#!/qPCRlibQnt). Libraries were diluted to 4nM, pooled, and denatured according to Illumina’s protocol. Balancing of pooled libraries was verified by sequencing on the MiSeq, using MiSeq Reagent Cartridge v2 300 cycles (Illumina, #15033624) and MiSeq Reagent Nano Kit v2 (Illumina, #15036714), at a concentration of 10pM. Libraries were sequenced by the Center for Genome Innovation at the University of Connecticut Institute for Systems Genomics on the Illumina NovaSeq 6000 at a concentration of 0.7nM.

For hESC-derived neurons, RNA libraries for RNA-seq were prepared using the TruSeq Stranded mRNA Library Prep Kit (Illumina, #20020594) following manufacturer’s protocols. Libraries were sequenced on the Illumina NextSeq 550 with settings for dual-index, paired-end sequencing, with 75 cycles per end at a concentration of 1.8pM.

### RNA-seq data processing

Quality control was performed on RNA-seq reads using FastQC (v.0.11.7) and MultiQC (v.1.10.1)(Ewels et al., 2016). Fastqs were aligned to hg38 using HISAT2 (v.2.2.1)(D. Kim et al., 2019)(Figure 2B & 3B)(Supplemental Tables 3 & 7), using options --fr --no-discordant --avoid-pseudogene --no-mixed, and STAR (v.2.7.1a)(Dobin et al., 2013)(Figure 1B-D; Figure 2A,C-F; Figure 3A, C-F)(Supplemental Tables 2 & 6), using options --readFilesCommand zcat --outFilterType BySJout --outFilterMultimapNmax 1 -- alignSJoverhangMin 8 --alignSJDBoverhangMin 2 --outFilterMismatchNoverReadLmax 0.04 -- alignIntronMin 20 --alignIntronMax 1000000 --alignMatesGapMax 1000000 --outSAMtype BAM SortedByCoordinate --outWigType bedGraph. Alignment to hg38 used gencode.v25.annotation.gtf. Equal distribution of reads across the gene body was verified using geneBody_coverage.py (v.3.0.1) from RSeQC (Wang et al., 2012). Sorted BAM files were used to extract read counts using featureCounts from subread (v.2.0)(Liao et al., 2014), with option -s 2.

### Differential gene expression (DEG) analysis

DEG analysis was performed in R (v.4.2.1)(R Core Team, 2022) on extracted read counts using DESeq2 (v.1.36.0)(Love et al., 2014). For comparisons of WT H9 ESCs to all neurons (Figure 1), low gene counts were filtered by removing all genes whose mean of counts across all samples was less than 39, or 1 count per sample. This resulted in a total of 25440 genes for downstream analysis. Pairwise differential analysis between WT ESCs and WT neurons, smDEL neurons, and lgDEL neurons was performed using the DESeqDataSetFromMatrix() with design = ∼ Condition_Lineage and a results() contrast of “Condition_Lineage”. For comparisons of lgDEL neruons to WT neurons (Figure 2) and smDEL neurons to WT neurons (Figure 3), unexpressed genes were removed from analysis by removing all genes whose mean of counts across all samples was less than 1. Three separate DESeqDataSetFromMatrix() designs were used for each lgDEL or smDEL comparisons (Figure 3E), ∼ Background_Condition, ∼ Condition, and ∼ Genetic.Background + Condition + Genetic.Background:Condition, with results() contrasts of “Background_Condition,” “Condition,” and “Condition,” respectively. For the lgDEL versus WT comparison, this resulted in 34921 genes for downstream analysis in the “Background_Condition” contrast and 33956 genes for downstream analysis in the “Condition” contrasts. For the smDEL versus WT comparison, this resulted in 33019 genes for downstream analysis in the “Background_Condition” contrast and 32715 genes for downstream analysis in the “Condition” contrasts. For all main figures, except for Figure 3F, the results from the “Background_Condition” contrast was used. For Figure 3F, the resultant genes from both lgDEL (691 genes) and smDEL (232 genes) comparisons were significant in all three analyses (Supplemental Figure 6B)(Supplemental Tables 2,6,10-13). PCA plots (Supplemental Figures 2A, 3A, 5A) were generated using the plotPCA() function from the DESeq2 package on rlog() transformed raw counts for filtered genes. To determine gene names from the ENSEMBL ID’s, biomaRt (v.2.52.0)(Durinck et al., 2005, 2009) was used with the ENSEMBL archive from April 2018 (host = https://apr2018.archive.ensembl.org/), the most similar database available for the GENCODEv25 annotation. Overlapping, or “shared,” genes were determined using ComplexUpset package (v.1.3.3)(Krassowski et al., 2021)(Figure 1C, 2A, 3A&F) in conjunction with ggplot2 (v.3.4.0)(Wickham, 2009) for graphing. Only significant DEGs were used for determining shared genes. To obtain significant DEGs, the default DESeq2 FDR setting (alpha) of 0.1 was used and then results tables (Supplemental Tables 2 & 6) were subsequently filtered for Benjamini-Hochberg (BH) adjusted p-values (padj) of < 0.05. Permutation tests were conducted to test the significance of these overlaps in the command line (Supplemental Figures 3B, 5B, 6C). Volcano plots were generated using the EnhancedVolcano package (v.1.14.0)(Blighe et al., 2018) with DEGs shared between H9 and CT2 backgrounds for both smDEL (Figure 2C-D) and lgDEL (Figure 3C-D); log2FoldChange for H9 and CT2 was averaged (x-axis).

### Gene and disease ontology analysis

Shared gene lists generated from DESeq2 analysis for either upregulated genes (Figure 1D), downregulated genes (Figure 2F), or upregulated and downregulated genes combined (Figure 2E, 3D) were processed for gene (GO) and disease ontology (DO) analysis using the clusterProfiler package (v.4.4.4)(Wu et al., 2021; Yu et al., 2012) and DOSE (v.3.22.1)(Yu et al., 2015) with functions of enrichGO() and enrichDGN() respectively with options for pAdjustMethod = “BH” and qvalueCutoff = 0.05. For enrichGO(), the org.Hs.eg.db package (v.3.15.0) was utilized. The universe used was the list of genes in each DESeq2 object generated via the “Background_Condition” contrasts (Figure 1D, 2E-F, 3D). GO results were simplified using the simplify() function from clusterProfiler with options of cutoff = 0.7 or 0.8. Dotplots were generated using ggplot2 on GO and DO results ordered first by the adjusted p-value then by foldEnrichment. The foldEnrichment score was calculated by dividing the *GeneRatio* by the *BgRatio* values for each result. For Figure 2E, a Gene-Concept Network plot was generated using the enrichplot package (v.1.16.2) using a foldChange object made with dplyr package (v.1.0.10). For Figure 4A, the disgenet2r package (v.0.99.3)(Piñero et al., 2020) was used, with a default universe of all genes.

### snoGloBe prediction and analysis

SnoGloBe was used to predict interactions of *SNORD116* C/D box snoRNAs with the 42 genes shared between small and large deletion models across both genetic backgrounds, as described previously (Deschamps-Francoeur et al., 2022). Per the authors’ recommendation to narrow the number of predictions obtained, we selectively kept the predicted interactions having at least three consecutive windows with a score of greater than or equal to 0.98 for further analysis, using options -t 0.98 -m -w 3. For the control analysis of 100 lists of 42 genes, we generated lists of genes which did not differ significantly from our list of 42 dysregulated genes (via the Wilcoxon test) in length, GC content, or expression in our inducible neuron system. These lists were then analyzed using snoGloBe for predicted binding of *SNORD116* using the same settings as above. Genomic feature coverage (Figure 4C)(Supplemental Figure 11) was determined using bedtools (v.2.29.0) for hg38. Only transcripts with support levels of 1-3 and a tag of basic were used. For plotting the distribution of the predicted region of interaction of snoRNAs (Figure 4D)(Supplemental Figure 12), the center of each binding event was determined and then the relative position of the binding event was calculated. The relative position of C/C’ and D/D’ boxes was calculated and then plotted using the grid.rect() function of the grid package (v.4.2.1).

## Data and code availability

Signal tracks for these experiments are available at the UCSC Genome Browser as a public session (https://genome.ucsc.edu/s/rbgilmore/PWS_RNAseq_bigwigs). All original code will be made publicly available on github (https://github.com/rachelgilmore/SNORD116_targets_functions.git). Upon publication, sequencing data will be available at the Gene Expression Omnibus (GEO). Cell lines are available upon reasonable request and after completion of Material Transfer Agreements through the University of Connecticut Cell and Genome Engineering Core. Any additional information required to reanalyze the data reported in this paper is available from the corresponding author upon request.

## Author Contributions

J.C., G.G.C., and R.B.G. conceived of and designed the study. Y.L., C.E.S., and M.S.C. provided resources. R.B.G. and Y.L. conducted investigation. R.B.G. performed formal analysis and generated figures. R.B.G. and J.C. wrote the original draft of the manuscript. All authors were involved in the editing and review of the manuscript, including approval of the final submitted version. J.C. and G.G.C. provided supervision. J.C. coordinated project administration.

## Acknowledgements

This work was supported by National Institutes of Health (NIH) grants T32HG010463 (R.B.G.), R35GM119465 (J.C.), and R01HD099975 (J.C. and G.G.C.). We thank the Center for Genome Innovation at the University of Connecticut Institute for Systems Genomics for their help in sequencing the ESC samples.

