## Supplemental_Figures for "Identifying key underlying regulatory networks and predicting targets of orphan C/D box *SNORD116* snoRNAs in Prader-Willi syndrome"

A)

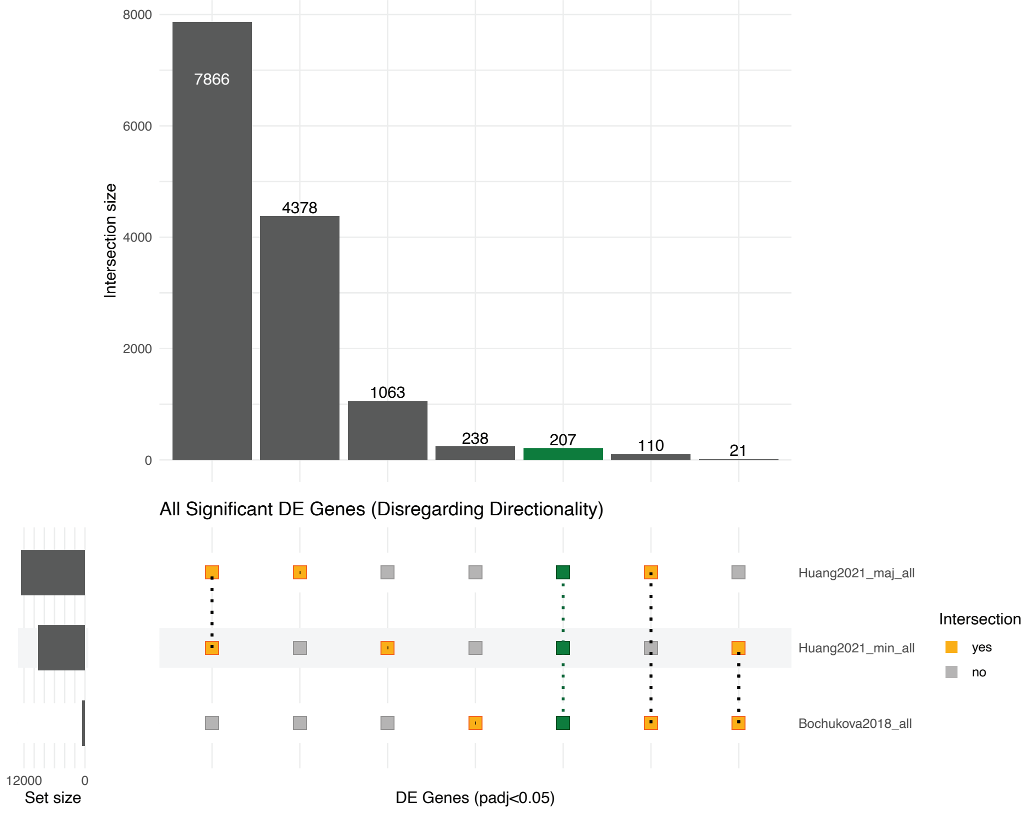


B)

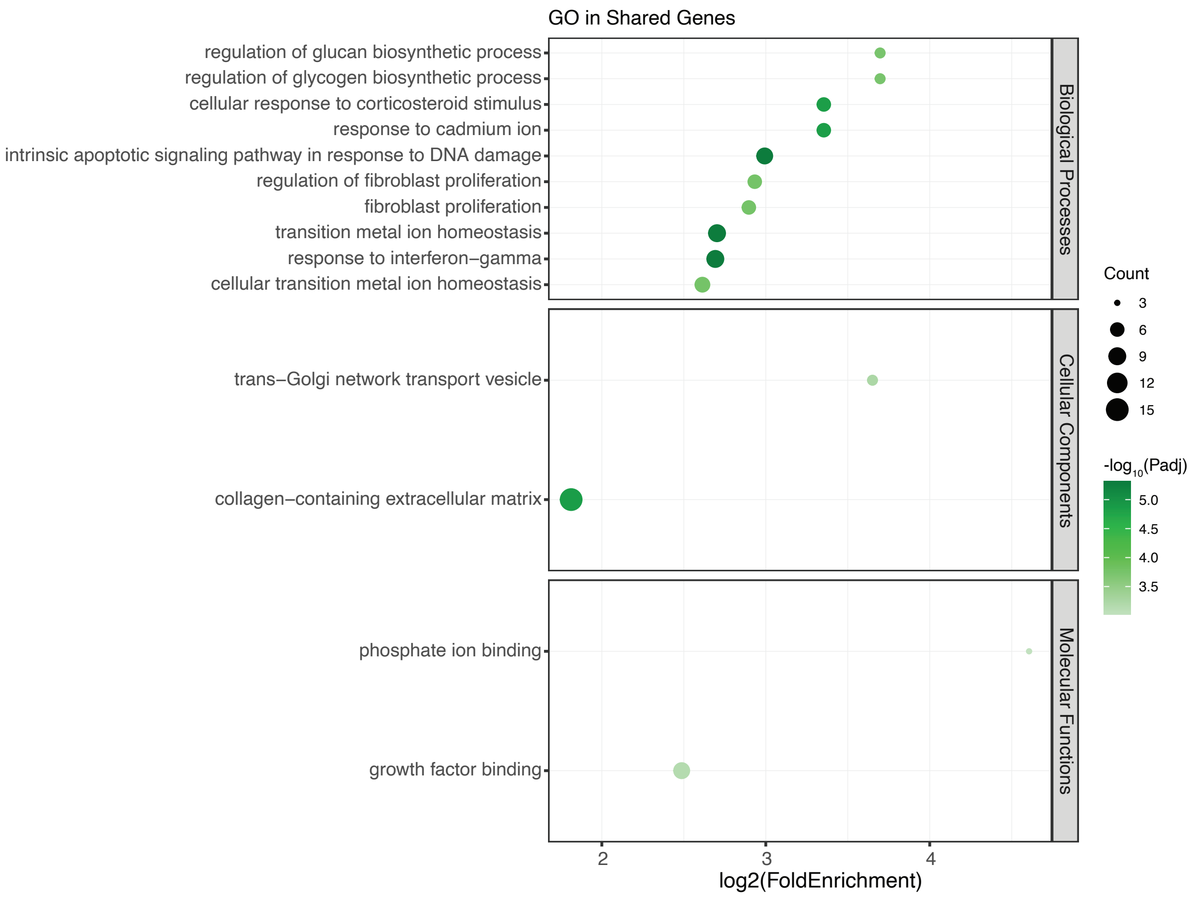


**Supplemental Figure 1. A)** Upset plot comparing significant DEGs (p.adjust < 0.05) in previously published studies. Green bar represents significant shared DEGs across all three data sets. **B)** Dot plot displaying gene ontology (GO) results for 207 shared dysregulated genes in all three studies. The x-axis represents the log2 fold enrichment value, and y-axis shows ontology terms. Size of the dot corresponds to the number of DEGs in our data set contained within each ontology term. Shading of the dot corresponds to the negative log10 of the adjusted p-value.

**A)**

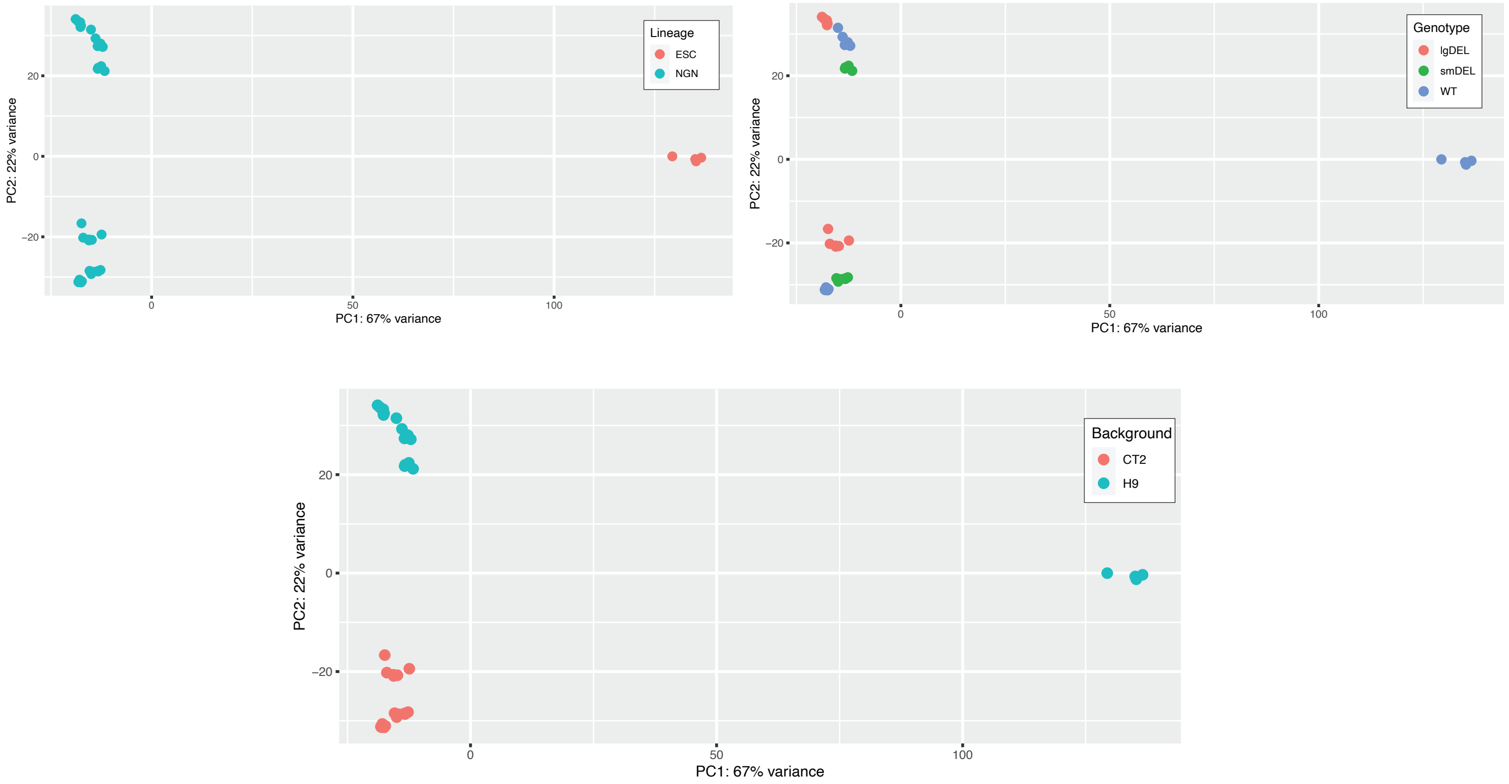


**B)**

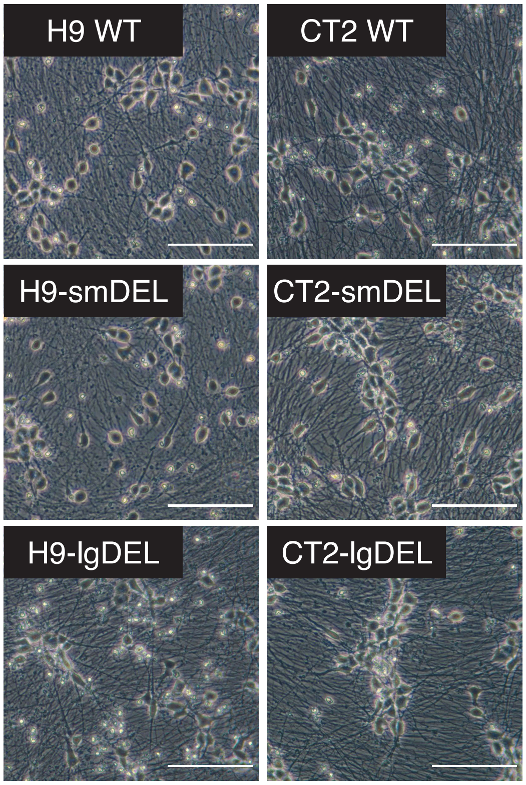


**C)**

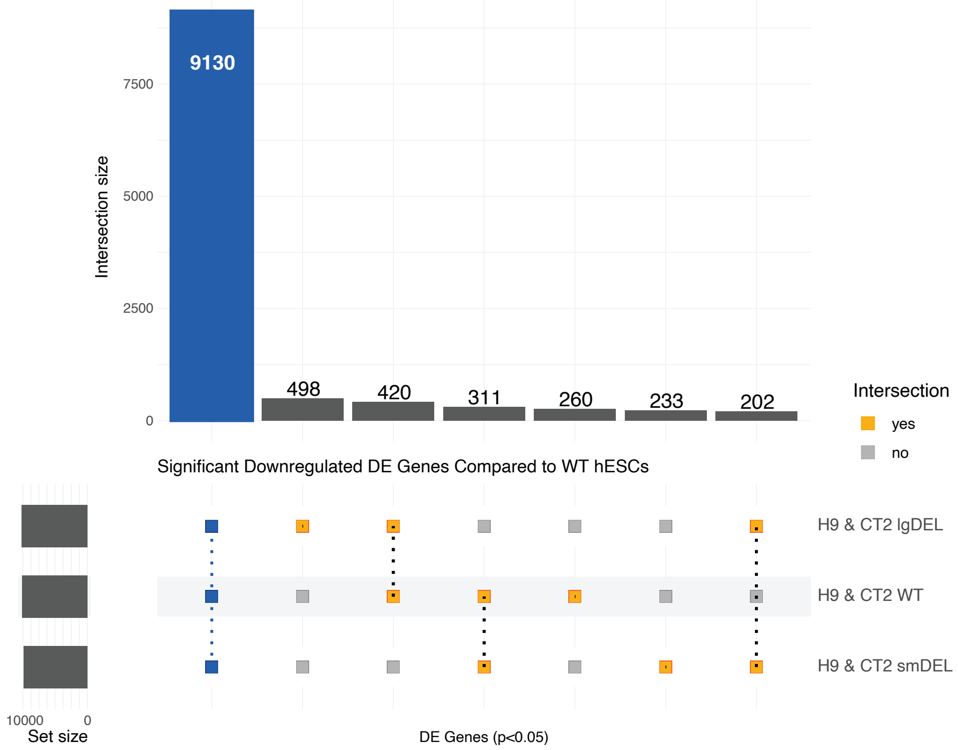


**Supplemental Figure 2. A)** PCA plots displaying variance of samples. Individual samples colored by lineage (*top left*), genotype (*top right*), or genetic background (*bottom middle*). **B)** Representative brightfield images of neurons taken at 20x magnification, scale bar is equal to 100 µm. **C)** Upset plot comparing significant DEGs (p.adjust < 0.05) of wild type (WT), smDEL, and lgDEL inducible neurons across both genetic backgrounds to wild type H9 ESCs. Blue bar represents significant shared downregulated ($log_2$FoldChange < 0) DEGs in all three genotypes (WT, smDEL, and lgDEL) versus WT ESCs.

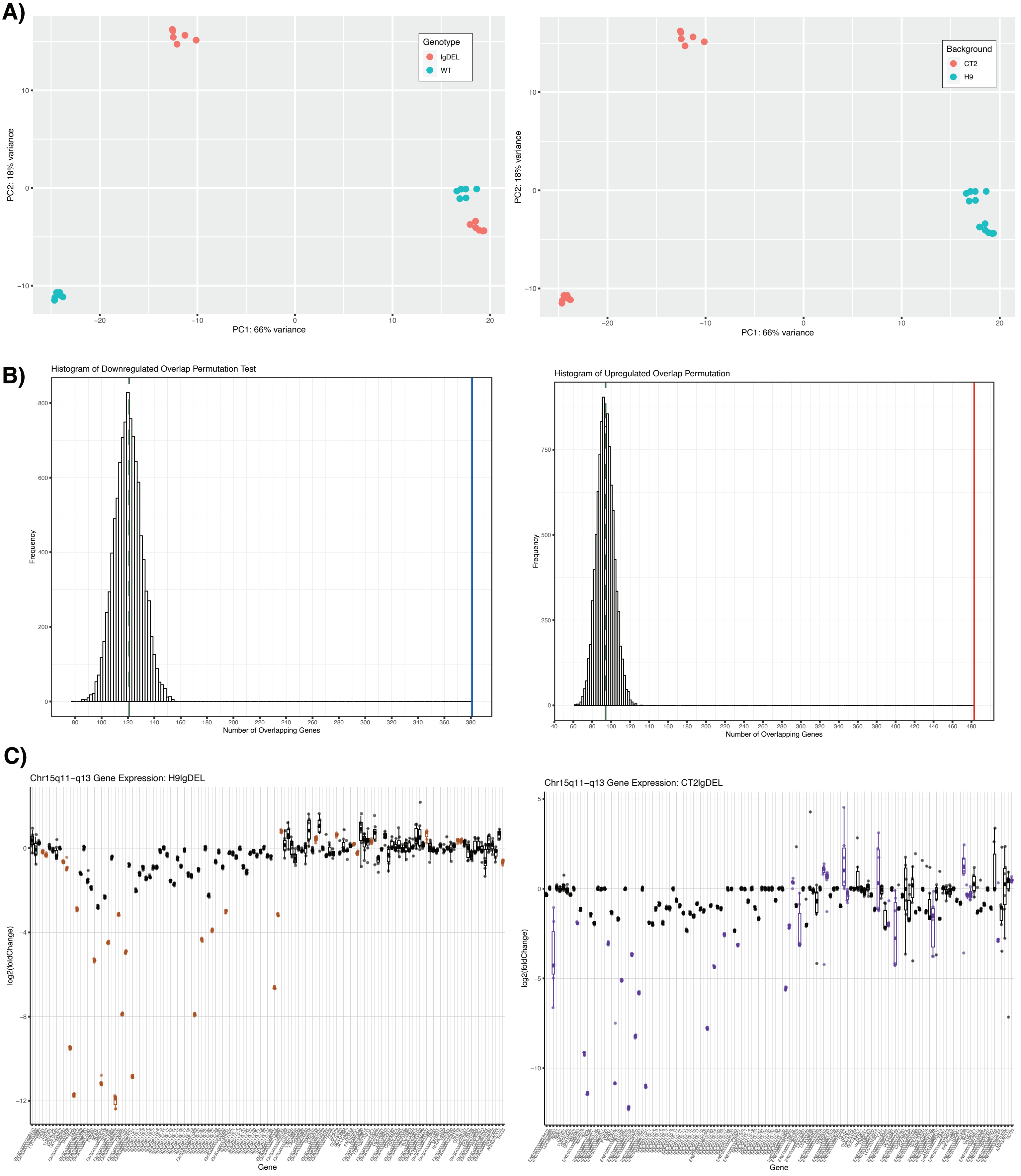


**Supplemental Figure 3. A)** PCA plots displaying variance of samples. Individual samples colored by genotype (*left*) or genetic background (*right*). **B)** Histogram of permutation test for overlapping genes. Green dashed line represents median number of overlaps. Solid blue bar (*left*) represents number of shared downregulated genes. Solid red line (*right*) represents number of shared upregulated genes. **C)** Box and whisker plots showing differential expression of chromosome 15q11-q13 region. Pseudocount was added to counts of all genes prior to calculation of log2(foldchange). Significant DEGs (p.adjust < 0.05) are shown in color, orange (H9lgDEL, *left*) or purple (CT2lgDEL, *right*).

A)

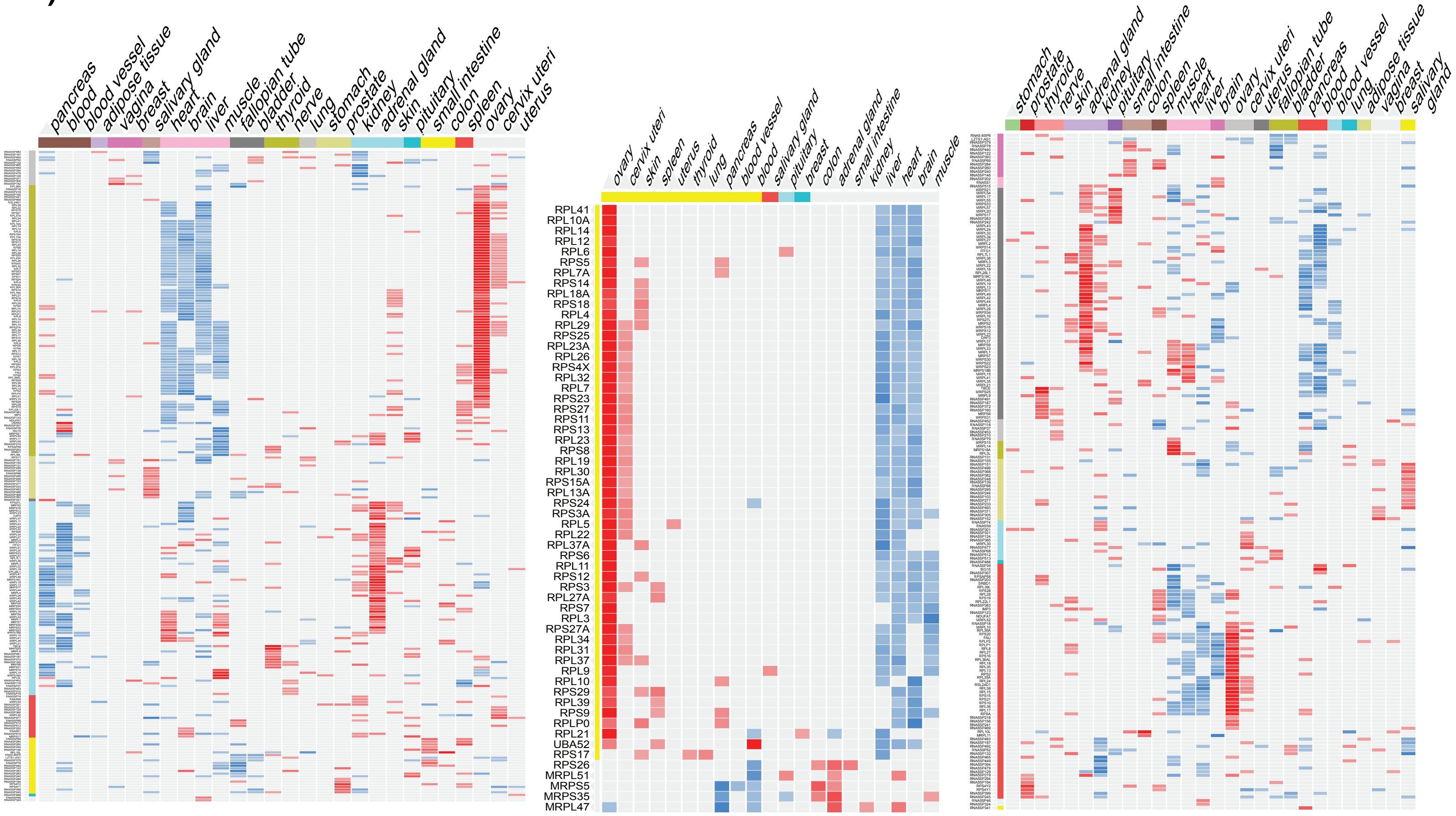


B)

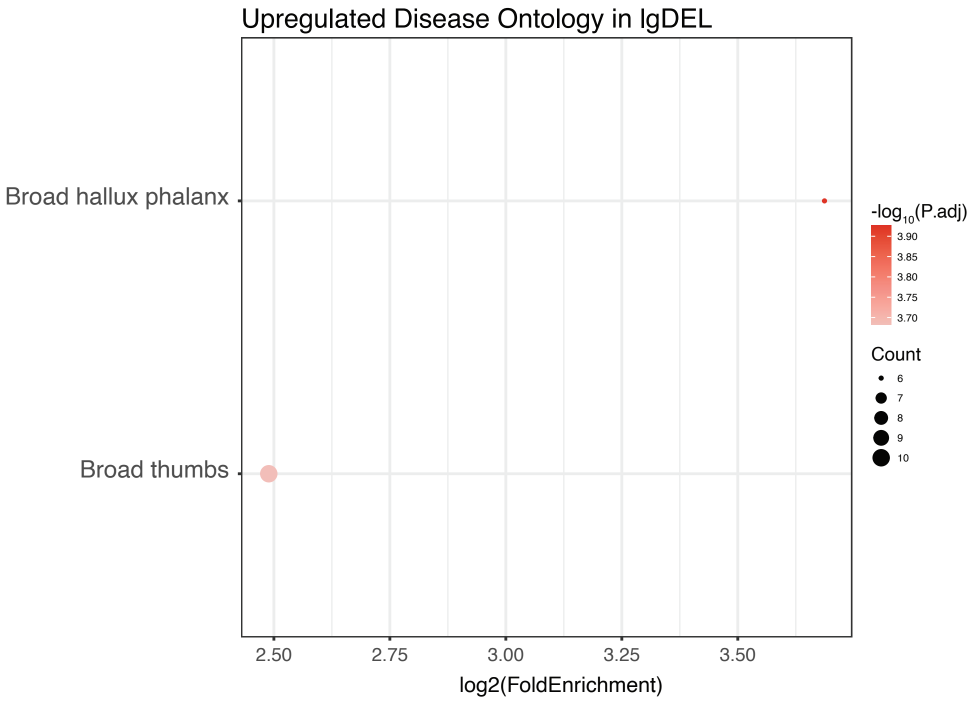


**Supplemental Figure 4. A)** Heatmaps generated using Harmonizome (v3.0, maayanlab.cloud/Harmonizome/visualize/heat_map/input_genes) for all genes contained within the structural constiuent of the ribosome ontology (GO0003735)(*left*), significant DEGs (*middle*), and all GO0003735 excluding the significant DEGs (*right*). The x-axis represents tissue types from which gene expression was profiled through GTEx, y-axis displays gene symbol. Row and column order were set to “Cluster”. Shading corresponds to gene expression, red being upregulated and blue being downregulated. **B)** Dot plot displaying disease ontology results for shared downregulated genes. The x-axis represents the log2 fold enrichment value, and y-axis shows disease ontology terms. Size of the dot corresponds to the number of DEGs in our data set contained within each ontology term. Shading of the dot corresponds to the negative log10 of the adjusted p-value.

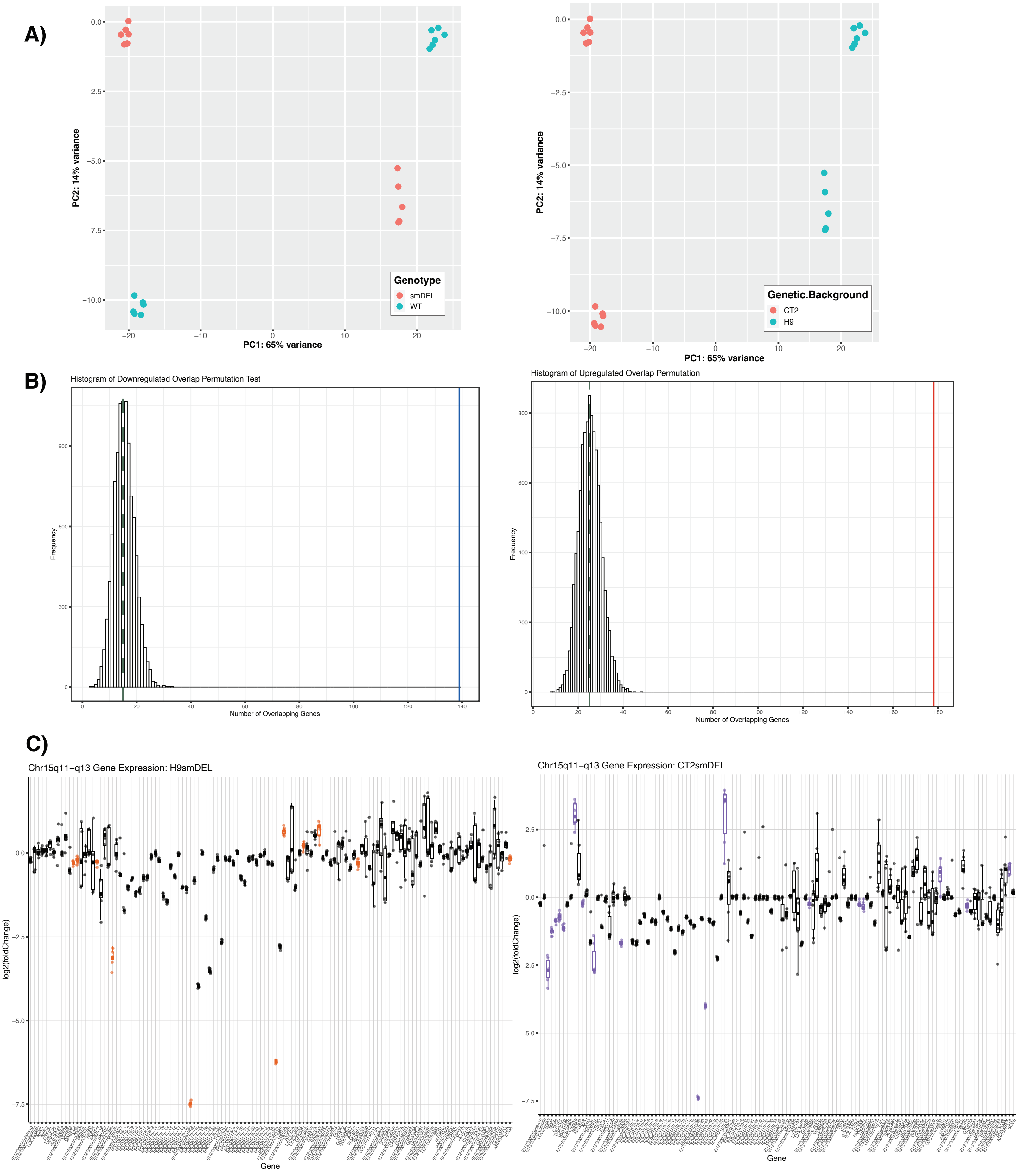


**Supplemental Figure 5. A)** PCA plots displaying variance of samples. Individual samples colored by genotype (*left*) or genetic background (*right*). **B)** Histogram of permutation test for overlapping genes. Green dashed line represents median number of overlaps. Solid blue bar (*left*) represents number of shared downregulated genes. Solid red line (*right*) represents number of shared upregulated genes. **C)** Box and whisker plots showing differential expression of chromosome 15q11-q13 region. Pseudocount was added to counts of all genes prior to calculation of log2(foldchange). Significant DEGs (p.adjust < 0.05) are shown in color, orange (H9smDEL, *left*) or purple (CT2smDEL, *right*).

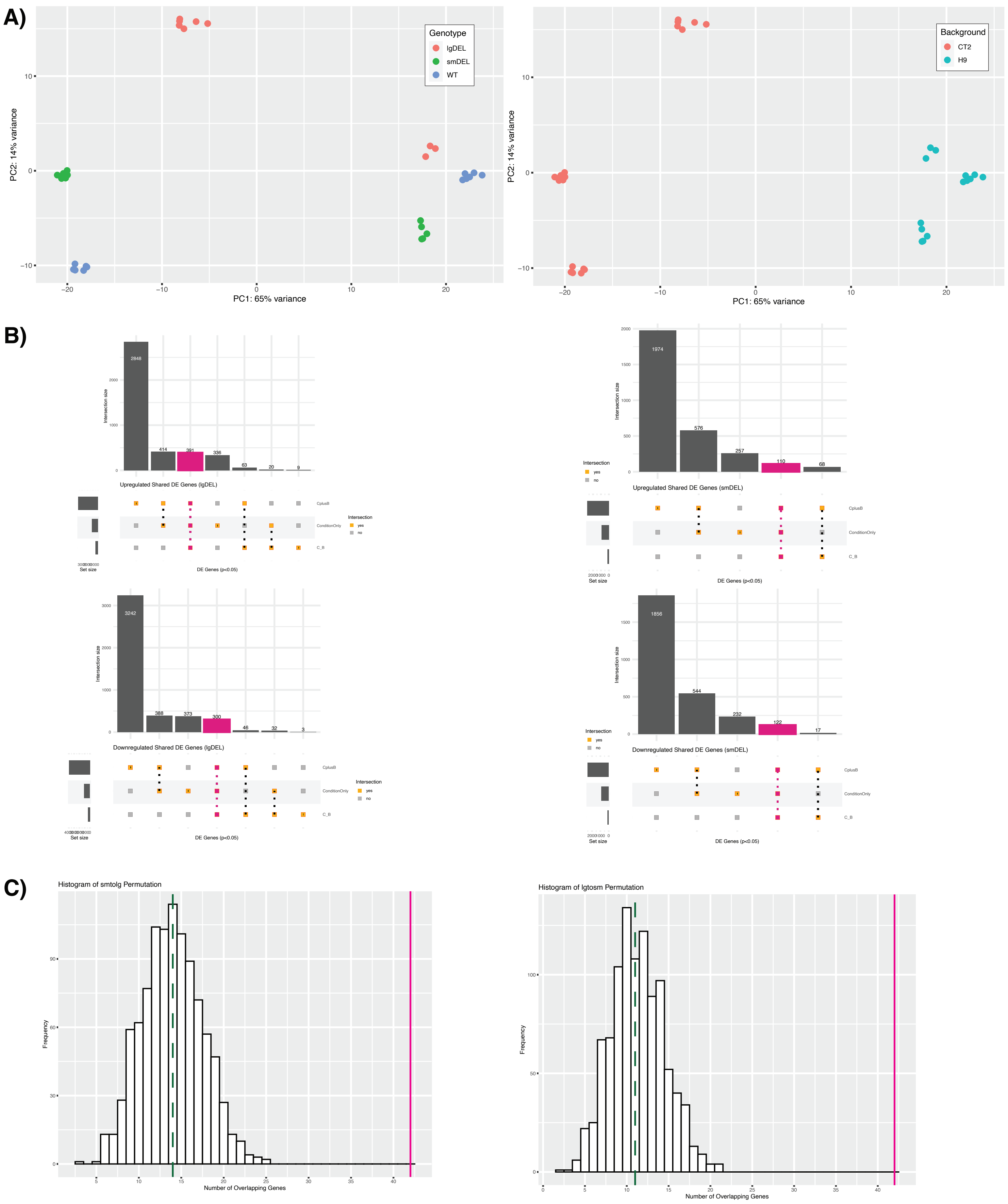


**Supplemental Figure 6. A)** PCA plots displaying variance of samples. Individual samples colored by genotype (*left*) or genetic background (*right*). **B)** Upset plot comparing significant DEGs (p.adjust < 0.05) of three separate DESeqDataSetFromMatrix() designs for lgDEL upregulated (top left, bottom left) and smDEL (top right, bottom right). Pink bars represent significant shared DEGs across each comparison. **C)** Histogram of permutation test for overlapping genes. Green dashed line represents median number of overlaps in either the permutation of overlaps in smDEL to lgDEL datasets (*left*) or lgDEL to smDEL datasets (*right*). Solid pink bar represents 42 shared dysregulated genes.

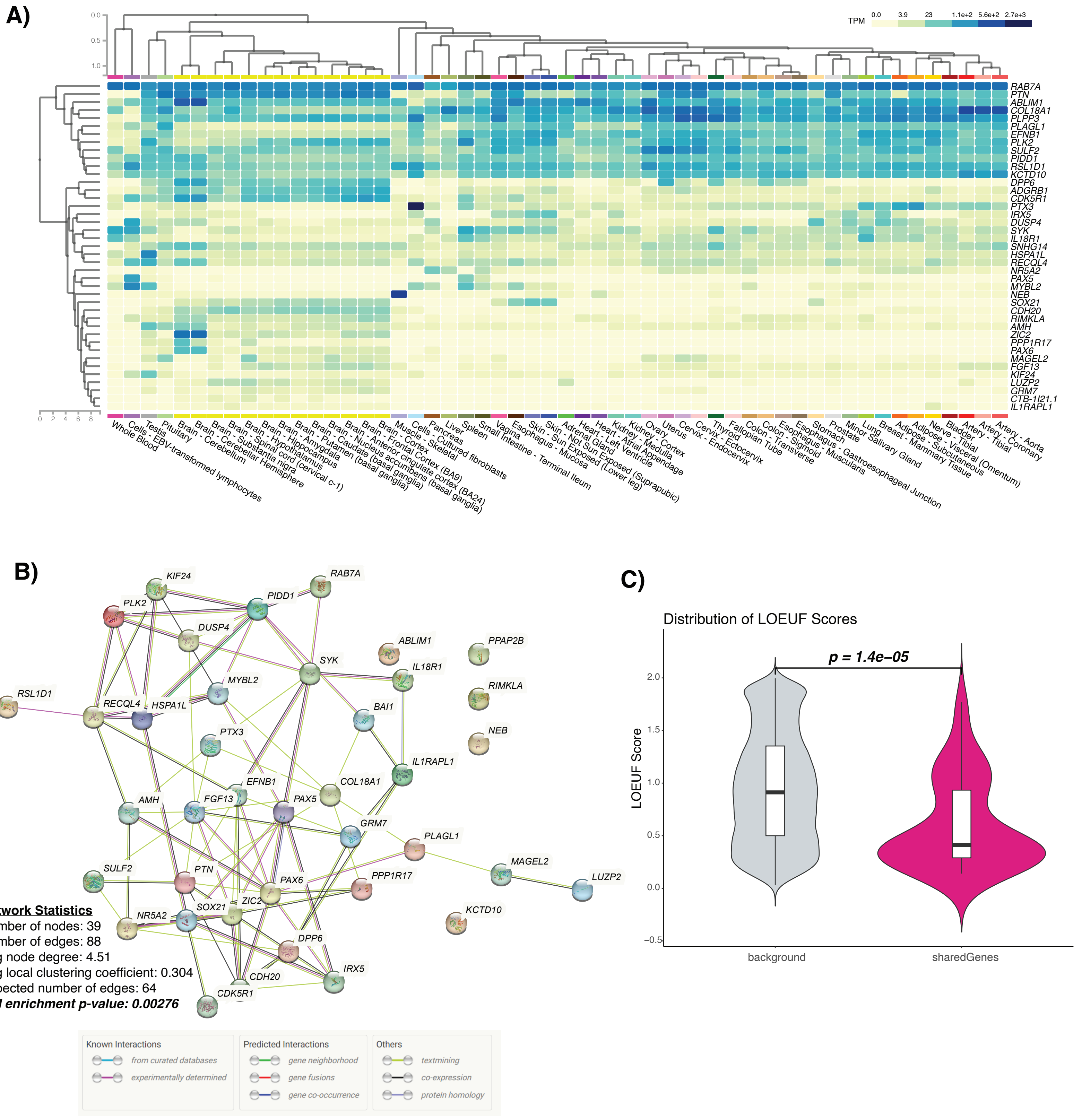


**Supplemental Figure 7. A)** Heatmap of expression of shared dysregulated genes from GTEx tissues. The x-axis displays the tissue type from which expression was profiled, and the y-axis displays the gene name. Shading of the graph corresponds to gene expression level in transcripts per million (TPM). Yellow indicates low level expression and dark blue indicates higher expression. **B)** Network of predicted protein interactions from our shared dysregulated genes generated using STRINGdb. Legend displaying colored interactions taken from string-db.org. **C)** Violin plot showing distribution of LOEUF scores of shared dysregulated genes (pink) compared to the rest of the gnomAD database (gray). Significance determined by Wilcoxon Rank Sum Test.

A)

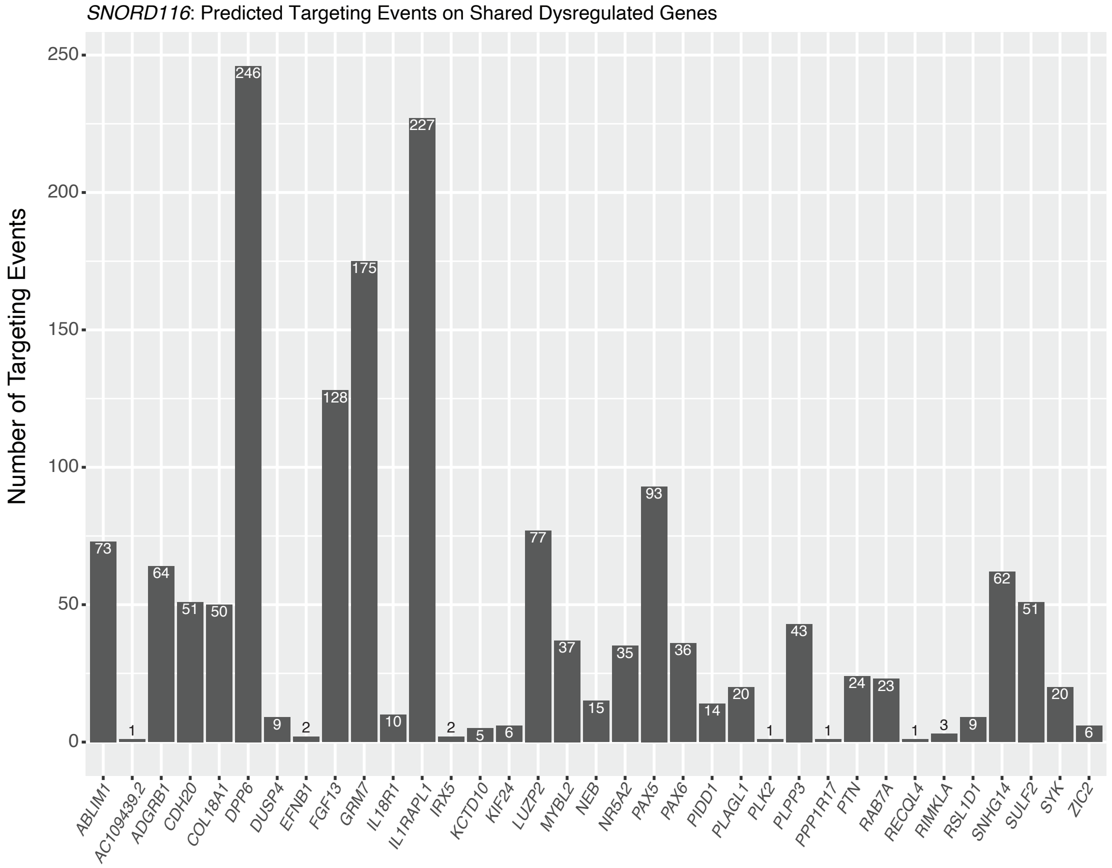


B)

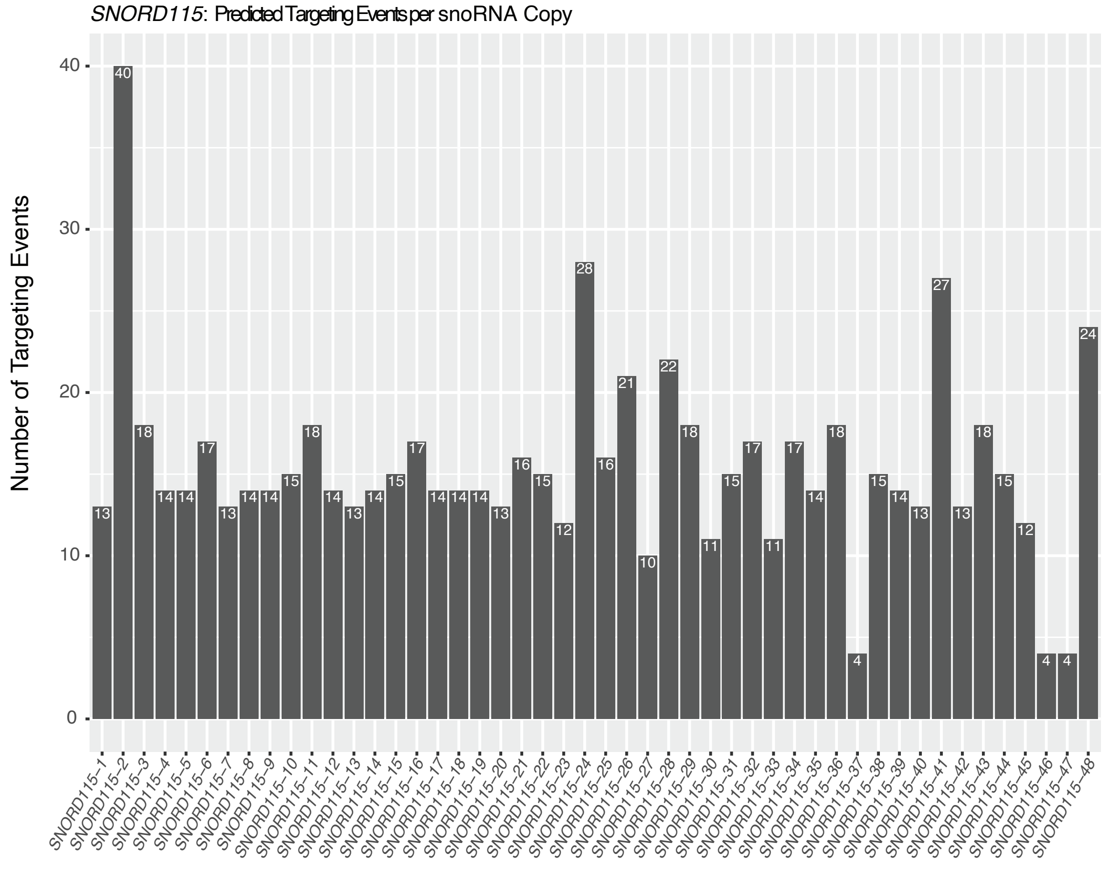


**Supplemental Figure 8. A)** Bar plot representing the number of predicted targeting events of *SNORD116* on shared list of dysregulated genes. The x-axis displays the predicted number of targeting events, and the y-axis displays the gene names. Note that this excludes predicted targeting events from three additional *SNORD116* copies found on chromosomes other than chr15. **B)** Bar plot representing the number of predicted targeting events per copy of *SNORD115*. The x-axis displays the predicted number of targeting events, and the y-axis displays the *SNORD115* copies. Note that this excludes two additional copies of *SNORD115* found on chromosomes other than chr15.

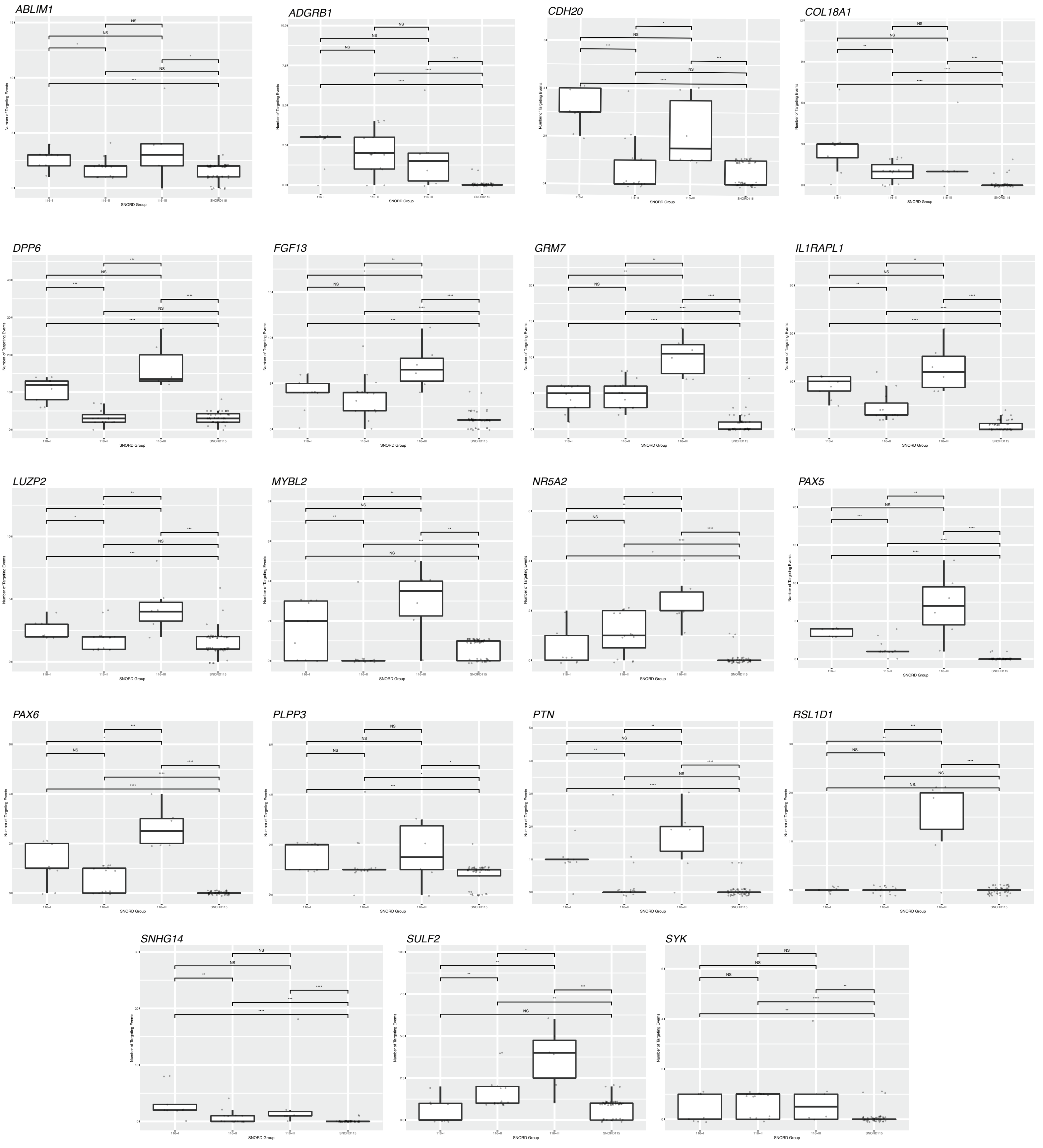


**Supplemental Figure 9.** Box plots displaying enrichment of *SNORD116-III* predicted targeting events versus *SNORD115*. The x-axis displays the SNORD group (*SNORD116-I*, *SNORD116-II*, *SNORD116-III*, and *SNORD115* from left to right). The y-axis represents the number of predicted targeting events. Each plot is an individual gene. Significance was determined by the Wilcoxon Test; NS = not significant (p-value > 0.05), * = p-value < 0.05, ** = p-value < 0.01, *** = p-value < 0.001, **** = p-value < 0.0001.

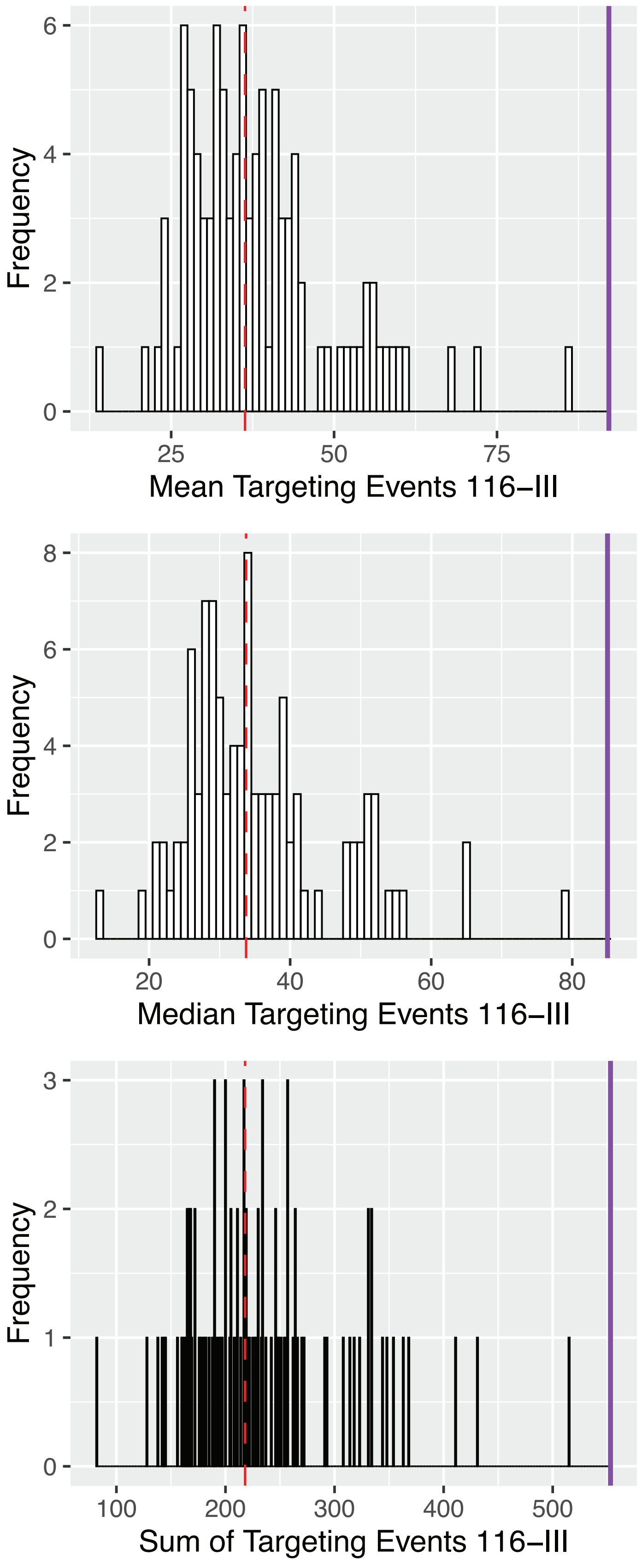


**Supplemental Figure 10.** Histogram of permutation test for predicted *SNORD116-III* targeting events on 100 random sets of 42 genes which did not differ significantly (via Wilcoxon test) from the set of 42 shared dysregulated genes in length, GC content, or expression in inducible neuron system. Red dashed line represents median number of mean (*top*), median (*middle*), or sum (*bottom*) of predicted targeting events. Solid purple bar represents number of experimentally obtained mean (*top*), median (*middle*), or sum (*bottom*) of predicted *SNORD116-III* targeting events on shared dysregulated genes, $p < 0.01$.

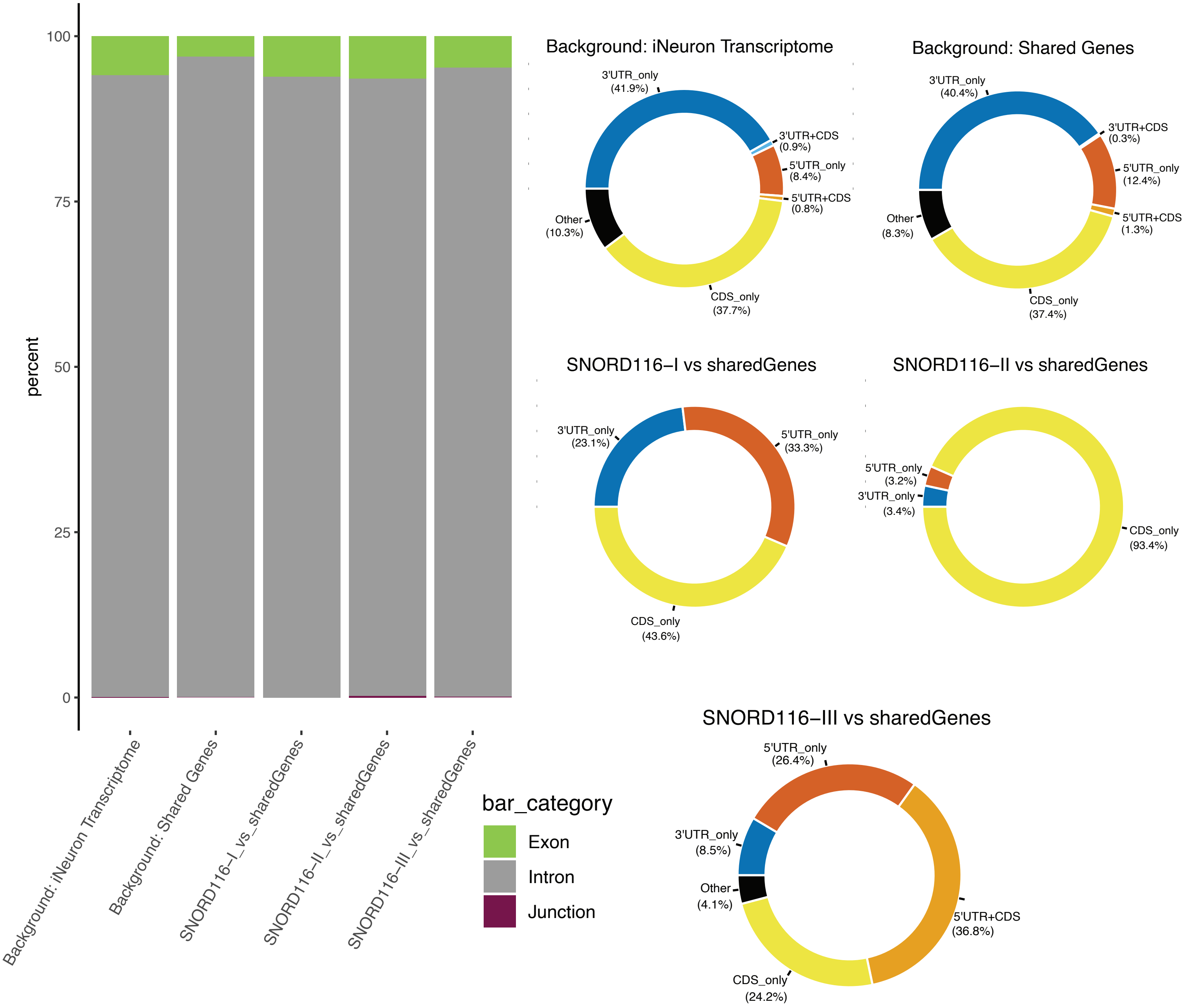


**Supplemental Figure 11.** Bar charts representing the proportion of exon, intron, and intron-exon junctions in the entire inducible neuron transcriptome, the set of 42 shared dysregulated genes, and the predicted targeting of *SNORD116-I*, *SNORD 116-II*, and *SNORD116-III* copies on those shared genes (*SNORD116-I/II/III* vs Shared Genes). Exon category is subdivided based on genic location and displayed as donut plots. Coloring of donut plots is based on exon category; 5'UTRs are represented in orange, 3'UTRs are represented in blue, CDS is represented in yellow, and any portion of exonic sequence not falling under those categories is termed "other" and shown in black.

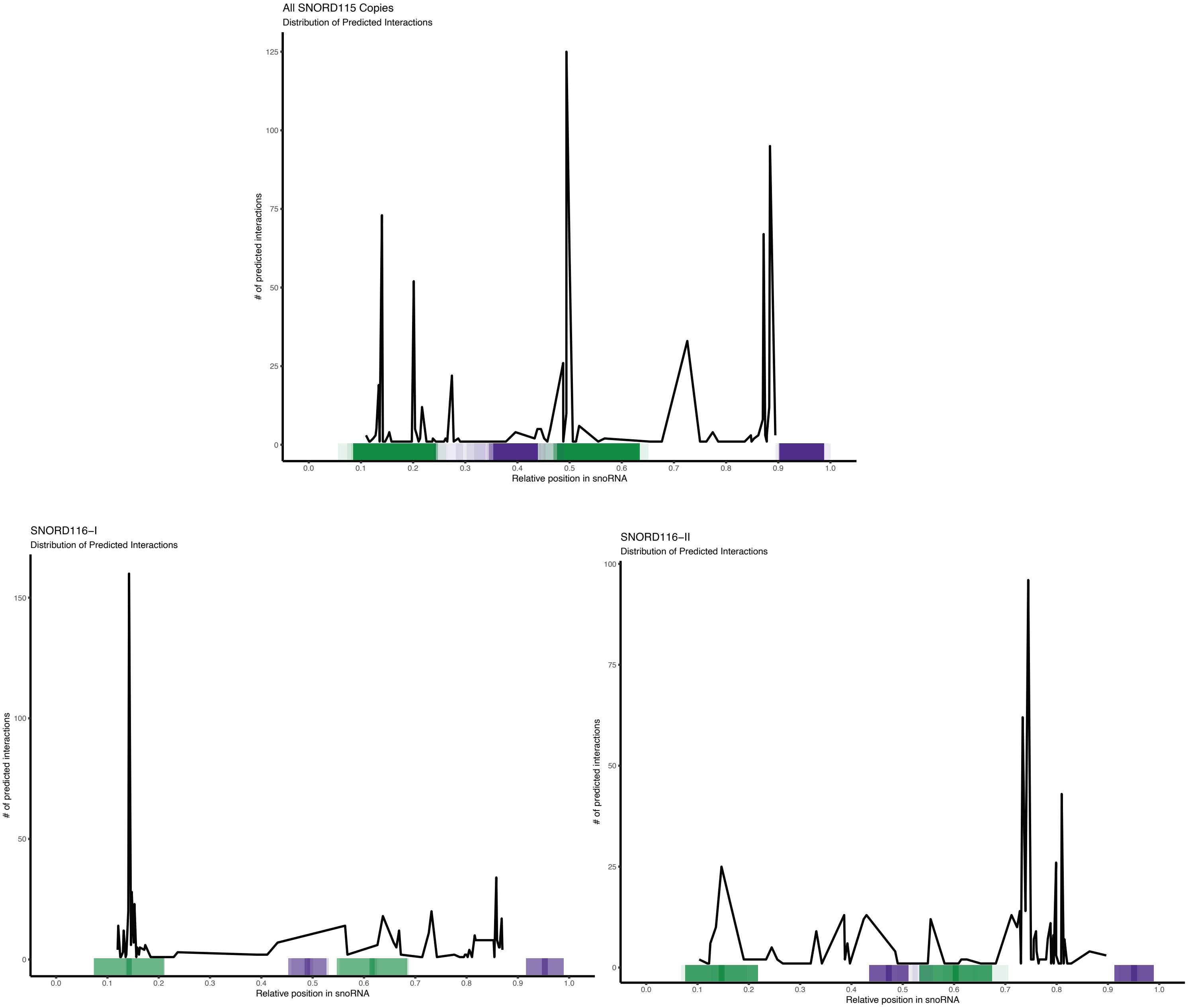


**Supplemental Figure 12.** Plots displaying distribution of prediction interactions for *SNORD115* (*top*), *SNORD116-I* (*bottom left*), and *SNORD116-II* (*bottom right*). The x-axis corresponds to the relative position within snoRNA copies, and y-axis represents the number of predicted interactions for which the center of the predicted binding interaction was used (black line). Color-coded bar on the x-axis indicates the position of C/C' and D/D' boxes found in snoRNA copies, indicated by green and purple respectively. The canonical antisense elements (ASEs) are found upstream of D/D' boxes.
